# Discovery and Optimization of Inhibitors for the Pup Proteasome System in *Mycobacterium tuberculosis*

**DOI:** 10.1101/796359

**Authors:** Guido V. Janssen, Susan Zhang, Remco Merkx, Christa Schiesswohl, Champak Chatterjee, K. Heran Darwin, Huib Ovaa

## Abstract

Tuberculosis is a global health problem with the existence and spreading of multidrug resistant and extensive drug resistant strains. The development of new drugs for tuberculosis that inhibit different activities than the current drugs is thus urgent. The prokaryotic ubiquitin like protein proteasome system is an attractive target for the development of new drugs. Using a Pup-based fluorogenic substrate, we screened for inhibitors of Dop, a depupylase, and identified I-OMe-Tyrphostin AG538 (**1**) and Tyrphostin AG53 (**2**). The hits were validated and determined to be fast reversible non-ATP competitive inhibitors. The SAR was established by testing 27 synthesized analogs of **1** and **2**. Several of the synthesized compounds also inhibited the depupylation of a native substrate, FabD∼Pup. Importantly, the pupylation and depupylation activities of PafA, the sole Pup ligase in *M. tuberculosis*, was also inhibited by some of these compounds. With the identification of the first described lead compounds for Dop and PafA inhibition, this study shows that high throughput screening can be a successful strategy for this purpose.

## Introduction

Tuberculosis (TB), the disease resulting from infection with *Mycobacterium tuberculosis* (*Mtb*) is one of the leading causes of death, killing almost two million people annually.^1^ The treatment for TB requires the use of one or more antibiotics that are taken daily for many months. This treatment can be accompanied by severe side effects like hepatitis, dyspepsia, exanthema and arthralgia and are a major factor to poor adherence in TB treatment.^2^ As a result, multidrug-resistant tuberculosis (MDR-TB) has developed that is resistant to both first line antibiotics isoniazid and rifampicin with or without the resistance to other drugs.^3,4^ More recently extensively drug-resistant tuberculosis and even totally drug-resistant tuberculosis have evolved of which the latter is resistant against all first- and second-line TB drugs.^5^ This poses a serious threat to human health and necessitates the development of new drugs to treat TB. These drugs should have novel modes of action, inhibiting targets different from those of currently used drugs, to circumvent resistances. Ideally, the targeted pathway should be specific for *Mtb* to minimize the possibility of unwanted side effects in the human host. An attractive target is the *Mtb* proteasome system.

The Pup-proteasome system (PPS) is essential for *Mtb* to cause lethal infections in animals.^6,7,8,9^ *Mtb* is one of the few bacterial orders that has proteasomes, which are large protein complexes that degrade proteins.^10^ The enzymatic activities of the proteasome core proteases are, for the most part, conserved between prokaryotes and eukaryotes.^11,12^ In general, proteins are degraded by a proteasome when they are post-translationally tagged with a small protein. In eukaryotes ubiquitin (Ub), a 76-amino acid small protein, serves as tag for substrate recognition by the proteasome.^13^ Ub is attached to a substrate *via* a multi-enzyme cascade consisting of E1, E2 and E3 enzymes, whereas the removal of Ub, is facilitated by a large family of deubiquitinating enzymes (DUBs).^14^ In *Mtb* a 64-amino acid prokaryotic ubiquitin-like protein (Pup), is responsible for substrate recognition by a bacterial proteasome. Compared to the ubiquitin proteasome system (UPS) where two E1, ∼40 E2, ∼600 E3 enzymes and ∼100 DUBs are involved, the Pup-proteasome system (PPS) is much less extensive; only two enzymes so far are needed for the pupylation and depupylation of substrates (Figure 1).

**Figure 1.**
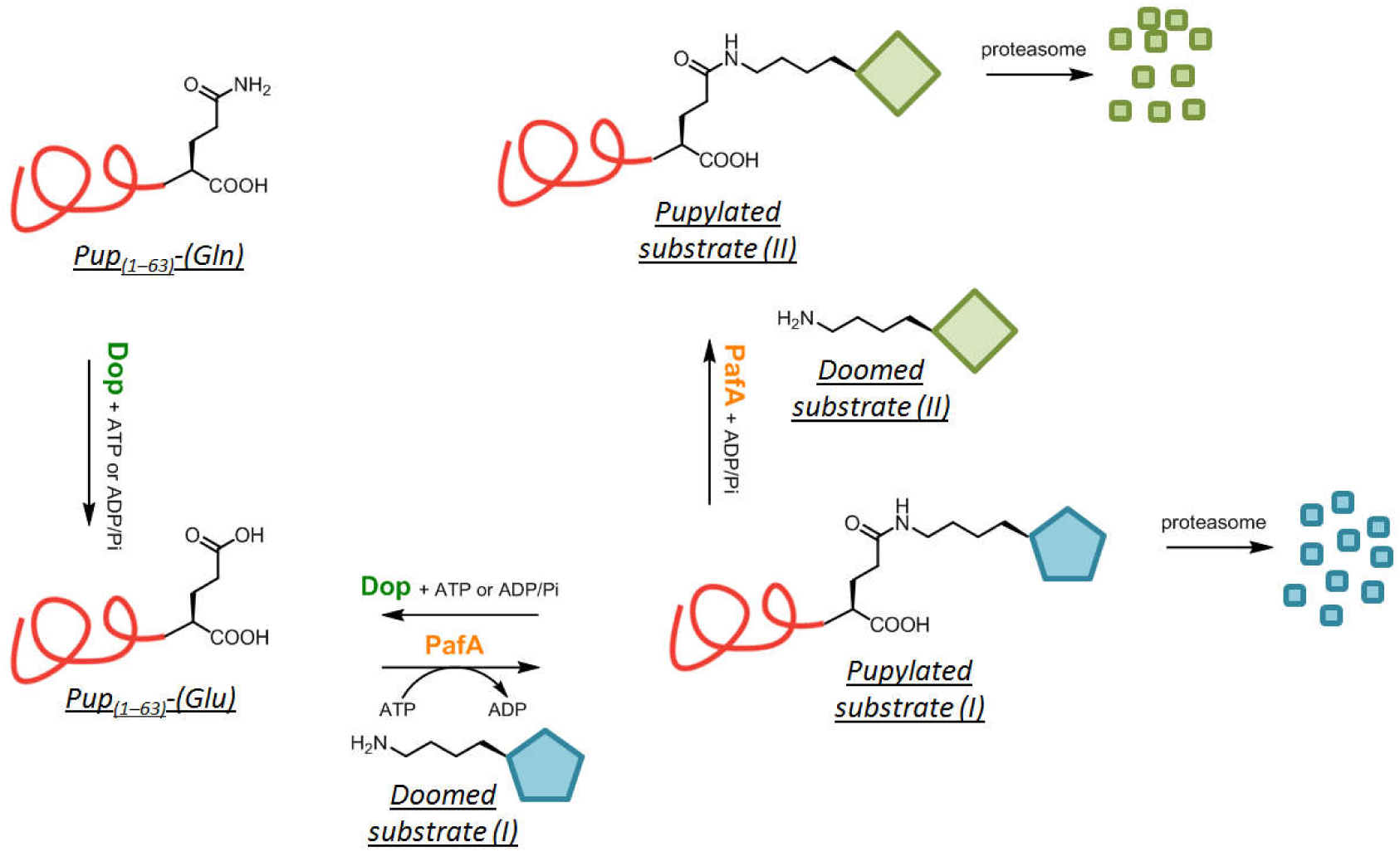
Schematic representation of the PPS in *Mtb*.

The sole ligase of Pup, PafA (proteasome accessory factor A), catalyzes the isopeptide bond formation between the γ-carboxylate of the C-terminal glutamate (Glu) of Pup with a lysine ε-amino group of a substrate. In this process, the C-terminal Glu is first phosphorylated by PafA, followed by nucleophilic attack of a substrate lysine ε-amino group to the activated carboxylic acid. As a result, PafA turns over ATP to ADP and Pi.^15^ In species where Pup is translated with a C-terminal glutamine (Gln), a deamidation step is needed to convert the terminal Gln to Glu by Dop (deamidase of Pup) before it can be ligated to a substrate by PafA.^15^ Although Dop and PafA have high sequence and structural similarity^15,16,17^ deamidation by Dop is not dependent on the hydrolysis of ATP.^18,19^

Both Dop^15,20^ and PafA^21,22^ can also depupylate substrates.^23^ Unlike pupylation, the precise mechanism of depupylation (and deamidation) is incompletely understood. While Dop also uses ATP as an activating co-factor, energy derived from its hydrolysis is not required^15,24,19^ The Weber-Ban group has suggested that ATP is hydrolyzed to ADP/Pi in order to assist in the hydrolysis of an amide bond.^25^ We have further proposed that a conserved aspartate (Asp) in Dop could act as a direct nucleophile to first attack an amide bond of Pup (either within Pup_Q_ or with a substrate) before hydrolysis.^19^ Relevantly this Asp residue is also conserved in PafA and is required for its pupylation and depupylation activities.^16,21^

Although Ub and Pup serve the same function, there is no sequence homology among the enzymes used for their conjugation or removal from substrates. Therefore, components of the PPS are attractive targets for the development of selective TB drugs with minimal side effects. Currently, only (4-(2-aminoethyl) benzenesulfonyl fluoride (AEBSF) is recently reported as inhibitor for the Pup ligase PafA and binds covalently to a serine (Ser119 in *Mtb* H37Rv PafA), thereby disturbing Pup recognition.^26^ AEBSF, however, is a pan-protease inhibitor that modifies active site serines.^13^ On the other hand, there are no known inhibitors for Dop.

In our efforts to find inhibitors to specifically target accessory factors of the *Mtb* PPS in the search for new drugs against TB, and to develop tools to study the mechanisms of action of these enzymes, we have previously developed a fluorogenic assay reagent to probe the activity of Dop *in vitro* that allows high throughput screening to identify inhibitors (Figure 2).^27^ The reagent is based on synthetically made, truncated Pup (Pup_33–63_) with a terminal Glu attached to 7-amino-4-methyl coumarin (AMC) *via* its γ-carboxylate. Cleavage of the AMC-group by Dop, but not PafA, releases AMC, which then fluoresces. Here we describe the use of this reagent in a small-molecule screen to identify inhibitors for Dop. Interestingly, several Dop-inhibitory compounds also inhibited both PafA activities. The hits were characterized in both biochemical and cell-based assays together with a structure activity relationship (SAR) study.

**Figure 2.**
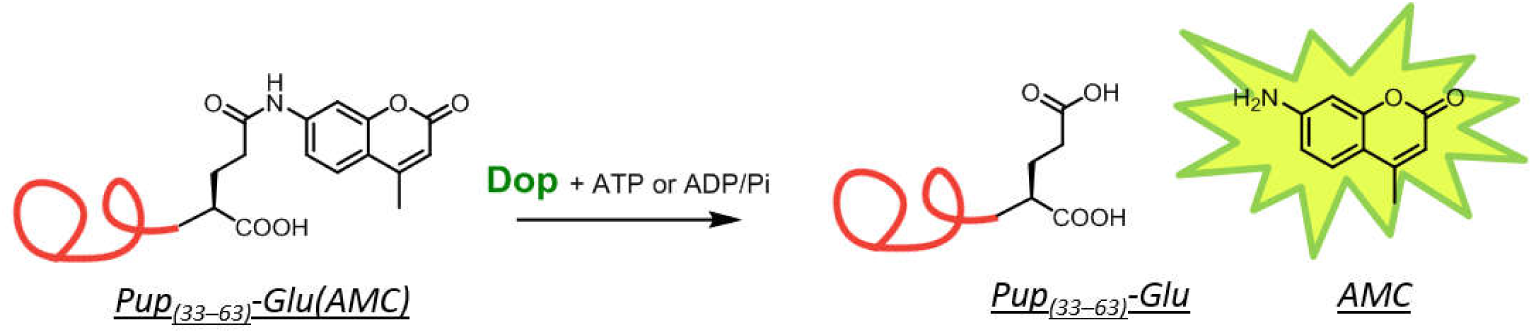
Fluorogenic assay to probe Dop activity, based on truncated Pup_(33–63)_-Glu and AMC.

## Results and Discussion

### Screen set up for Dop inhibitors

We screened a library of 1280-pharmaceutically active compounds (Lopac) for Dop inhibitors using our previously developed fluorogenic Dop substrate, Pup_(33–63)_-Glu(AMC).^27^ We identified several hits in this screen, including I-OMe-Tyrphostin AG 538 (**1**) and Tyrphostin AG538 (**2**), which were the most potent compounds (Figure 2). Both molecules consist of a *cis*-benzylidenemalonitrile backbone that is on both sides connected to aromatic ringsCompounds **1** and **2** have been identified before as inhibitors for several kinases with various modes of inhibition. For example, **1** and **2** act as substrate compettive inhibitors of the IGF-1 Receptor Kinase^28^, while they inhibit phosphoinotiside kinase PI5P4Kα by competing with ATP.^29^ On the other hand, oxidative stress-sensitive Ca^2+^-preamble channel TRPA1 and TPRM2 activated by H_2_O_2_, are inhibited by scavenging hydroxyl radicals.^30,31^

To validate the hits, we repeated the inhibition experiment with **1** and **2** determined the IC_50_ values (Figure 3). Values of 0.52±0.12 and 0.3±0.03 μM were found for **1** and **2** respectively. We next assessed the specificity of **1**, by performing a ‘ratio test’ (Supplementary information Figure 1).^32^ In this assay, the IC_50_ value of **1** was measured at a routine and 10-fold increased Dop concentration. No significant change was found in IC_50_ values for **1** (IC_50_ = 0.26±0.06 μM for 30 nM Dop; IC_50_ = 0.70±0.03 μM for 300 nM Dop), showing that **1** does not inhibit Dop *via* aggregation.

**Figure 3.**
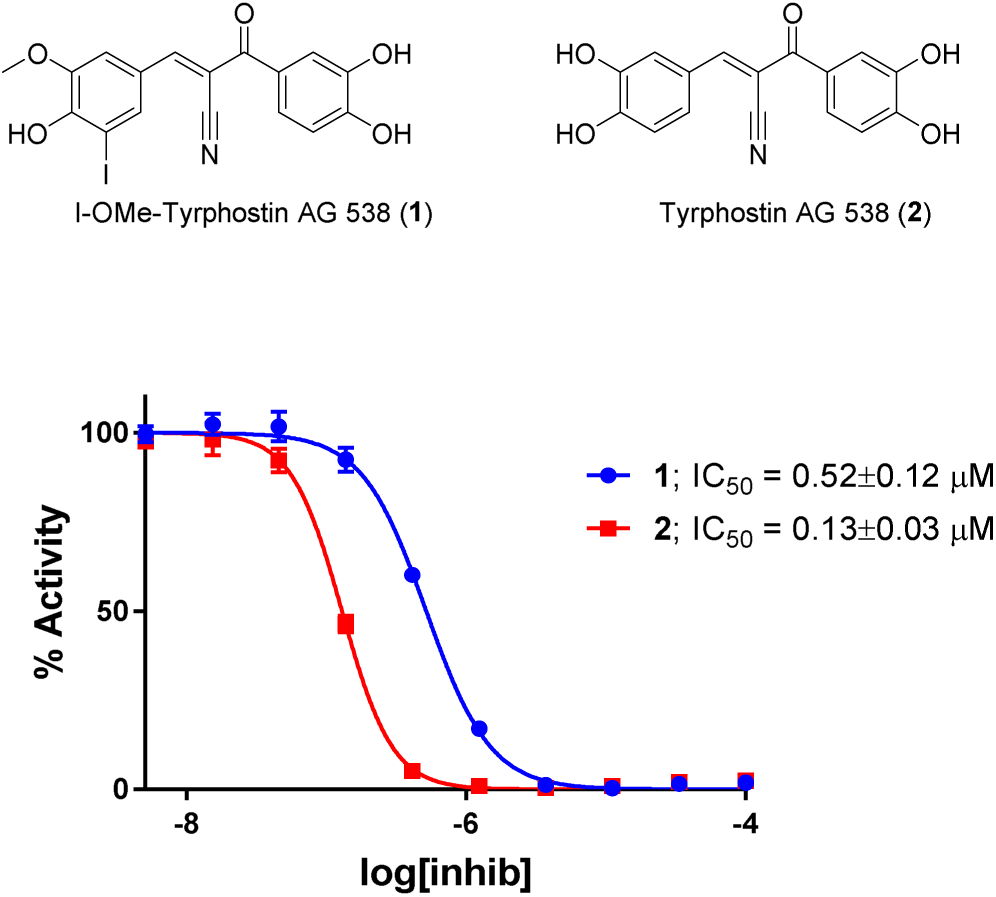
Chemical structures and IC_50_ curves for **1** and **2**.

To get an understanding of the mode of Dop inhibition by **1**, we first tested if different incubation times of **1** together with Dop would result in a change of IC_50_ value. If **1** were irreversibly and covalently inhibiting Dop, lower IC_50_ values would be expected with longer incubation times, because **1** has more time to form a covalent bond with Dop. Thus, the IC_50_ values were determined with 5 and 60 minutes pre-incubation of Dop with **1** before addition of the substrate (Supplementary Figure 2). The IC_50_ values did not change significantly (0.47±0.05 μM for 5 min incubation *vs.* 0.71±0.08 μM for 60 min incubation) indicating a non-covalent mode of inhibition. This was further confirmed by the results of a ‘jump-dilution’ assay (Supplementary information Figure S3).^33,34^ In this assay Dop was pre-incubated for 30 minutes with a concentration of 10 times the IC_50_ value for **1**, followed by a 100-fold dilution and reassessment of the activity. For **1**, complete restoration of Dop activity was observed after dilution, suggesting that there was a fast re-equilibration of the Dop-**1** complex and that **1** is a fast-reversible inhibitor.

Next, we tested if **1** inhibited Dop by competing with the binding of ATP. Therefore, we first determined the K_m_ of ATP for Dop (0.32±0.03 mM, Figure 4A). Subsequently, the IC_50_ value was determined in the presence of eight different concentrations around the K_m_ of ATP ranging from 0.1–6.4 mM (Figure 4B). In the case of an ATP competitive inhibitor, a positive relationship between the IC_50_ and log[ATP] was expected.^34,29^ Because the IC_50_ did not significantly increase with higher ATP concentration, we concluded that **1** is not an ATP competitive inhibitor.

**Figure 4.**
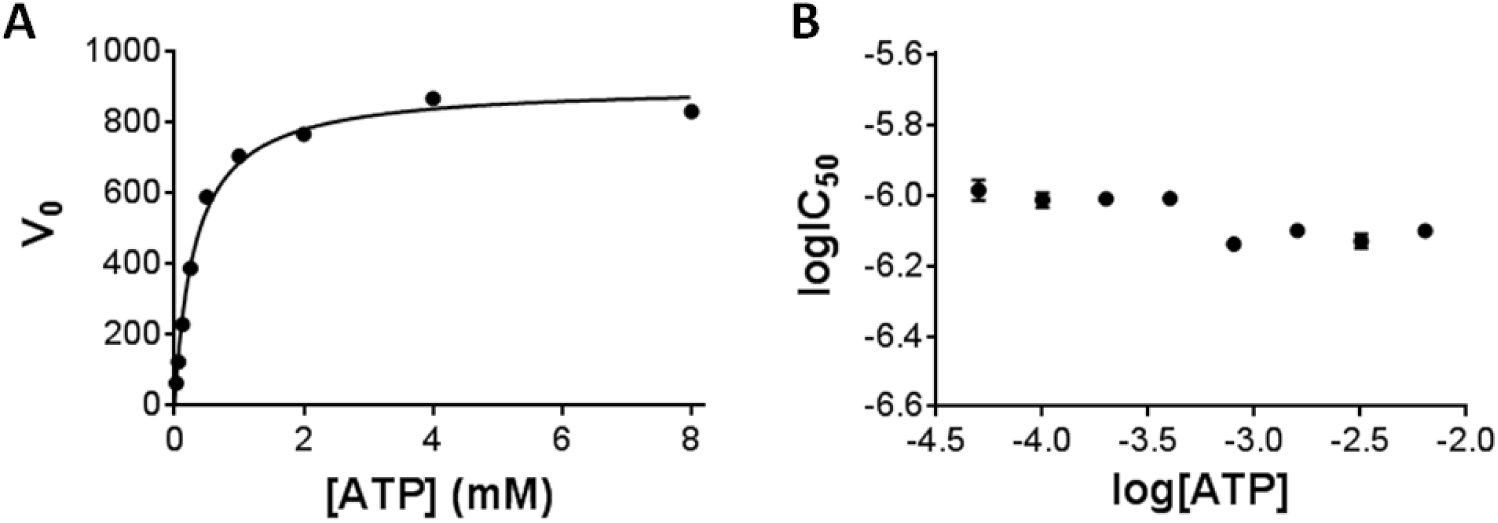
(A) Michaelis-Menten curve of Dop *vs.* [ATP]. K_m_ = 0.32 ± 0.03 mM, V*max* = 907 ± 22. (B) ATP competition assay with **1**: the IC_50_ of **1** for Dop is plotted against the log[ATP]. Eight ATP concentrations ranging from 0.3–20 times the K_m_ of ATP were evaluated in duplicate.

With **1** validated as a specific, fast reversible and non-ATP competitive inhibitor for Dop, we used it as lead compound in the subsequent SAR study focusing on variation of substituents on the aromatic rings followed by modification of the benzilydenemalonitrile core (Figure 5). In this work, a total number of 27 compounds were evaluated for their potency to inhibit the activity of Dop using the Pup_(33–63)_-Glu(AMC) assay by measuring the IC_50_ values (Table 1).

**Figure 5.**
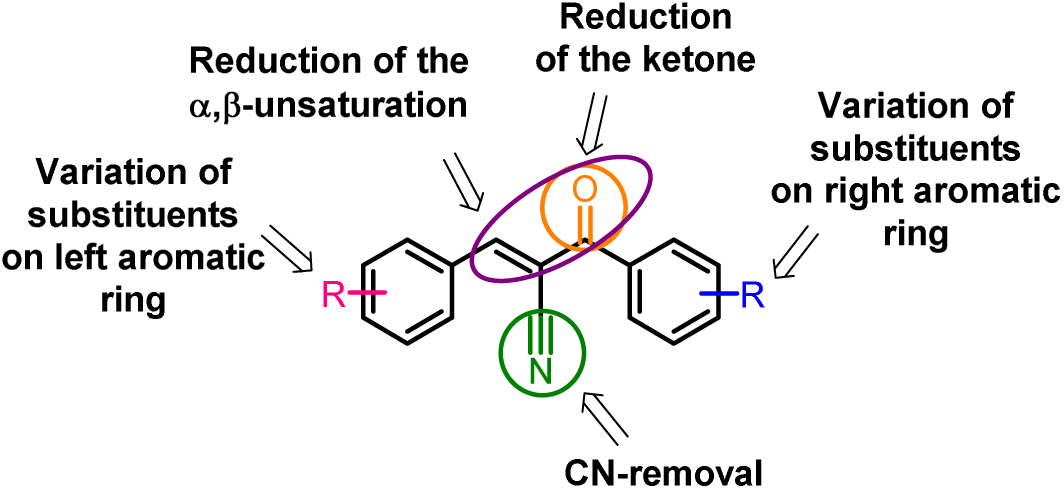
Focus areas for the SAR study.

**Table 1.**
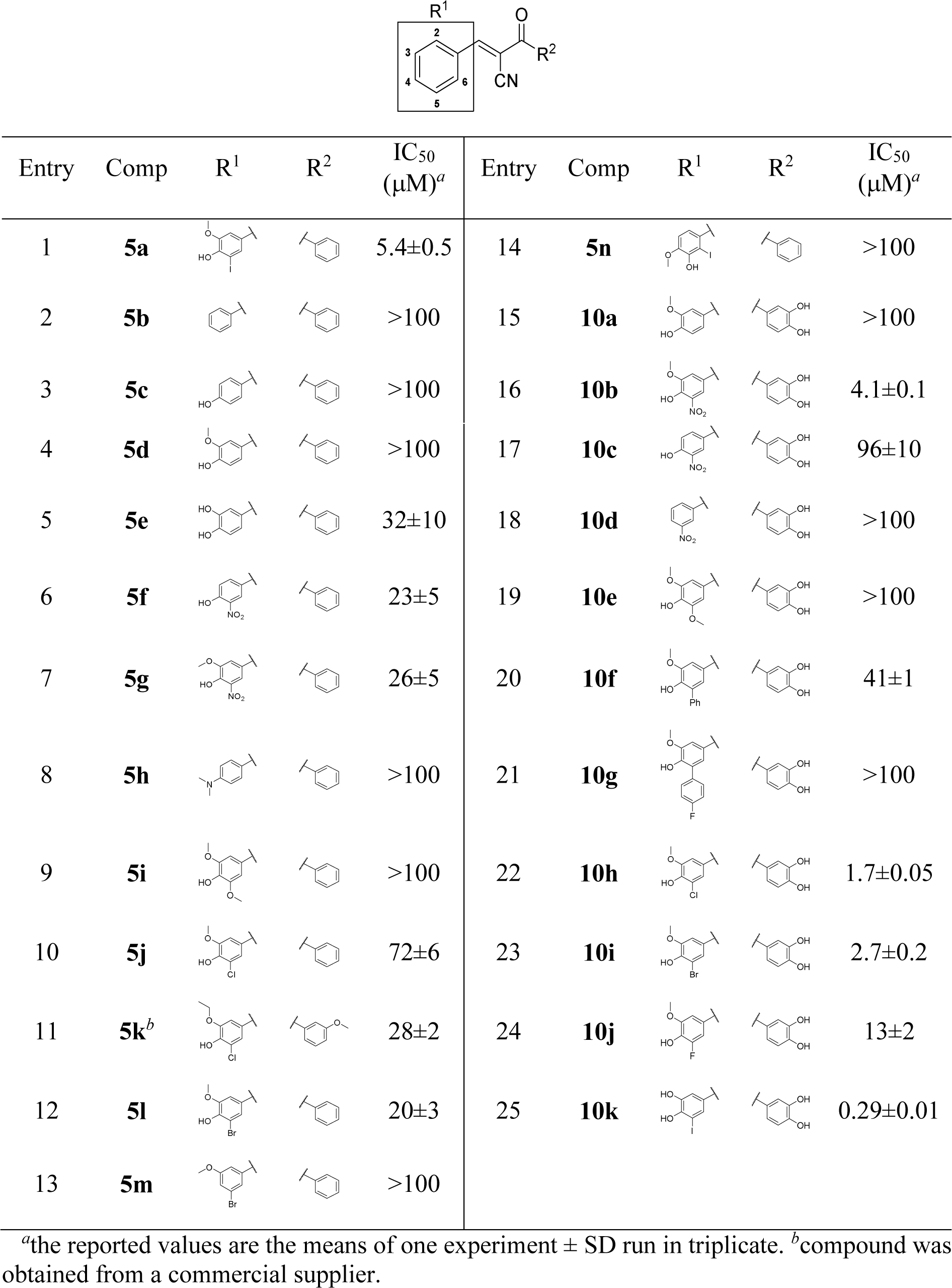
Inhibitory activity of the synthesized compounds for Dop.

|  |  |  |  |  |  |  |  |  |  |
| --- | --- | --- | --- | --- | --- | --- | --- | --- | --- |
| Entry | Comp | R<sup>1</sup> | R<sup>2</sup> | IC<sub>50</sub><br>(μM)<sup>a</sup> | Entry | Comp | R<sup>1</sup> | R<sup>2</sup> | IC<sub>50</sub><br>(μM)<sup>a</sup> |
| 1 | **5a** | | | 5.4±0.5 | 14 | **5n** | | | >100 |
| 2 | **5b** |  |  | >100 | 15 | **10a** |  |  | >100 |
| 3 | **5c** | | | >100 | 16 | **10b** | | | 4.1±0.1 |
| 4 | **5d** |  |  | >100 | 17 | **10c** |  |  | 96±10 |
| 5 | **5e** |  |  | 32±10 | 18 | **10d** |  |  | >100 |
| 6 | **5f** |  |  | 23±5 | 19 | **10e** |  |  | >100 |
| 7 | **5g** |  |  | 26±5 | 20 | **10f** |  |  | 41±1 |
| 8 | **5h** |  |  | >100 | 21 | **10g** |  |  | >100 |
| 9 | **5i** |  |  | >100 | 22 | **10h** |  |  | 1.7±0.05 |
| 10 | **5j** |  |  | 72±6 | 23 | **10i** |  |  | 2.7±0.2 |
| 11 | **5k<sup>b</sup>** |  |  | 28±2 | 24 | **10j** |  |  | 13±2 |
| 12 | **5l** |  |  | 20±3 | 25 | **10k** |  |  | 0.29±0.01 |
| 13 | **5m** |  |  | >100 |  |  |  |  |  |

<sup>a</sup>the reported values are the means of one experiment ± SD run in triplicate. <sup>b</sup>compound was obtained from a commercial supplier.

### Chemistry

The target compounds were close derivatives of **1** and **2** (Table 1). The benzylidenemalonitrile core (**5**, Scheme 1) was conveniently prepared through a Knoevenagel condensation reported by Kaufmann^35^ involving treatment of benzaldehydes (**3**) and malonitrile (**4**) with piperidine in ethanol to give the desired compounds in 35–85% yield (Scheme 1).

For the synthesis of compounds **10a**–**10m**, bearing two hydroxyl groups on the malonitrile part, we envisioned 3,4-dimethoxymethyl malonitrile **8** as the reaction partner for the Knoevenagel reaction. Compound **8** was synthesized from 3,4-dihydroxybenzoic acid methyl ester that was efficiently MOM-protected using K_2_CO_3_ in acetone to afford **6** in quantitative yield. Subsequent substitution of the ester **7** using deprotonated acetonitrile afforded malonitrile **8** in 95% yield. The Knoevenagel reaction using **8** proceeds smoothly with a range of benzaldehydes. Deprotection of the hydroxyl groups is conveniently achieved using a mild method that employs KHSO_4_ coated silicagel^36^ to afford the final compounds **10a**–**10m** in good yield after purification.

Compound **12** that is lacking the nitrile moiety on the Michael acceptor was synthesized from aldehyde **3a** and acetophenone (**11**) using KOH as base in a similar Knoevenagel condensation (Scheme 2A). Complete reduction of the Michael acceptor to alcohol **13** was achieved in 22% yield as a 2:1 mixture of diastereomers by treating **5a** with NaBH_4_ in methanol for 15 minutes (Scheme 2B). Additionally, we selectively reduced the carbonyl of **5a** by applying Luche’s reduction conditions (NaBH_4_/CeCl_3_) to obtain **14** in 50% yield (Scheme 2C). Unfortunately, our attempts to selectively reduce the double bond while maintaining the carbonyl moiety were unsuccessful.

**Scheme 1.**
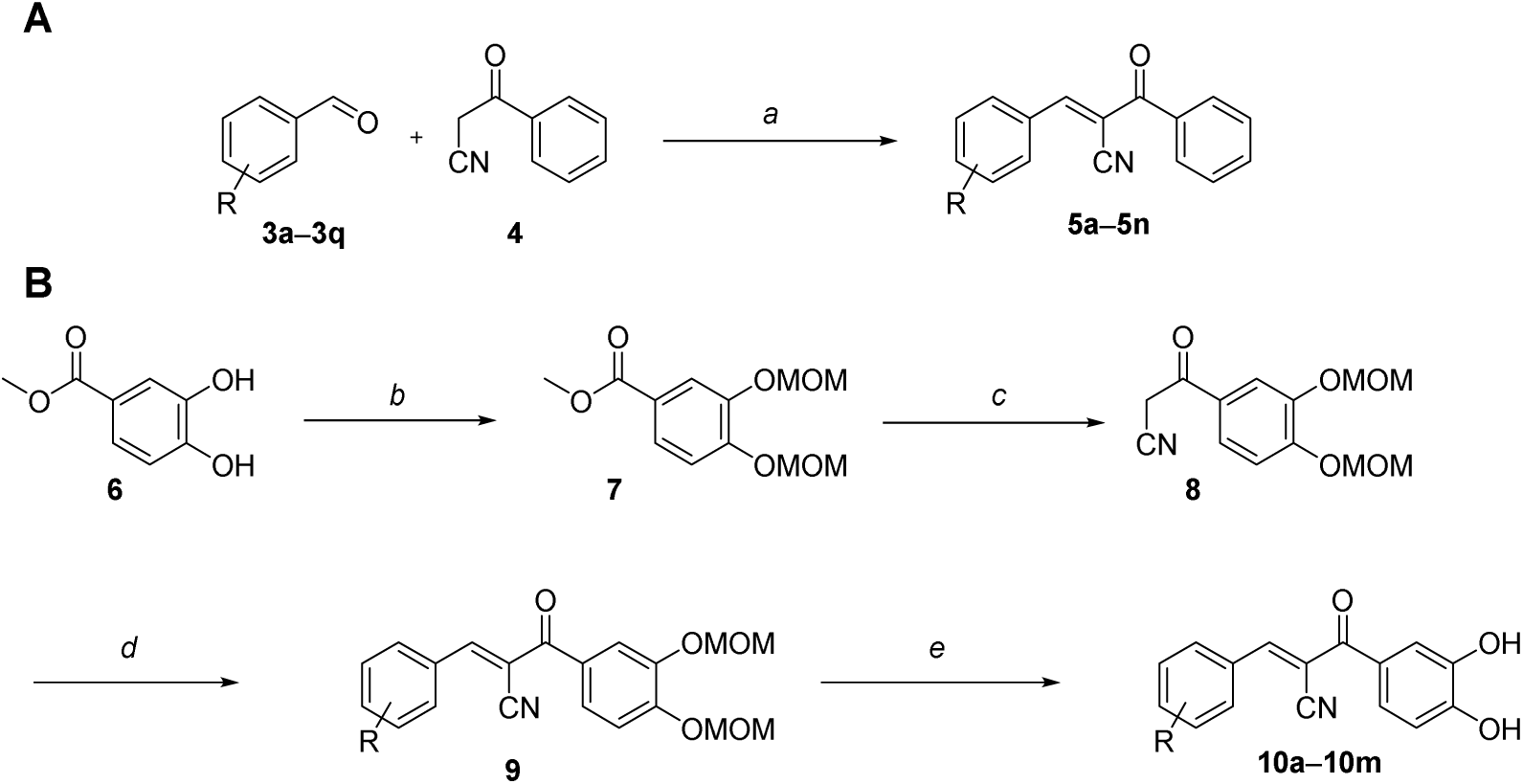
Synthetic routes for the synthesis of compounds **5** and **10**^*a*^. ^*a*^Reagents and conditions: (a) piperidine, EtOH, room temperature 16h (35–85%); (b) MOMCl, K_2_CO_3_, acetone 60 °C overnight (95%); (c) LiCH_2_CN, THF, –78 °C, 2h (95%); (d) ArCHO, piperidine, EtOH, 20 °C overnight; (e) KHSO_4_·SiO_2_, CH_2_Cl_2_, r.t. overnight (20–77% over two steps).

**Scheme 2.**
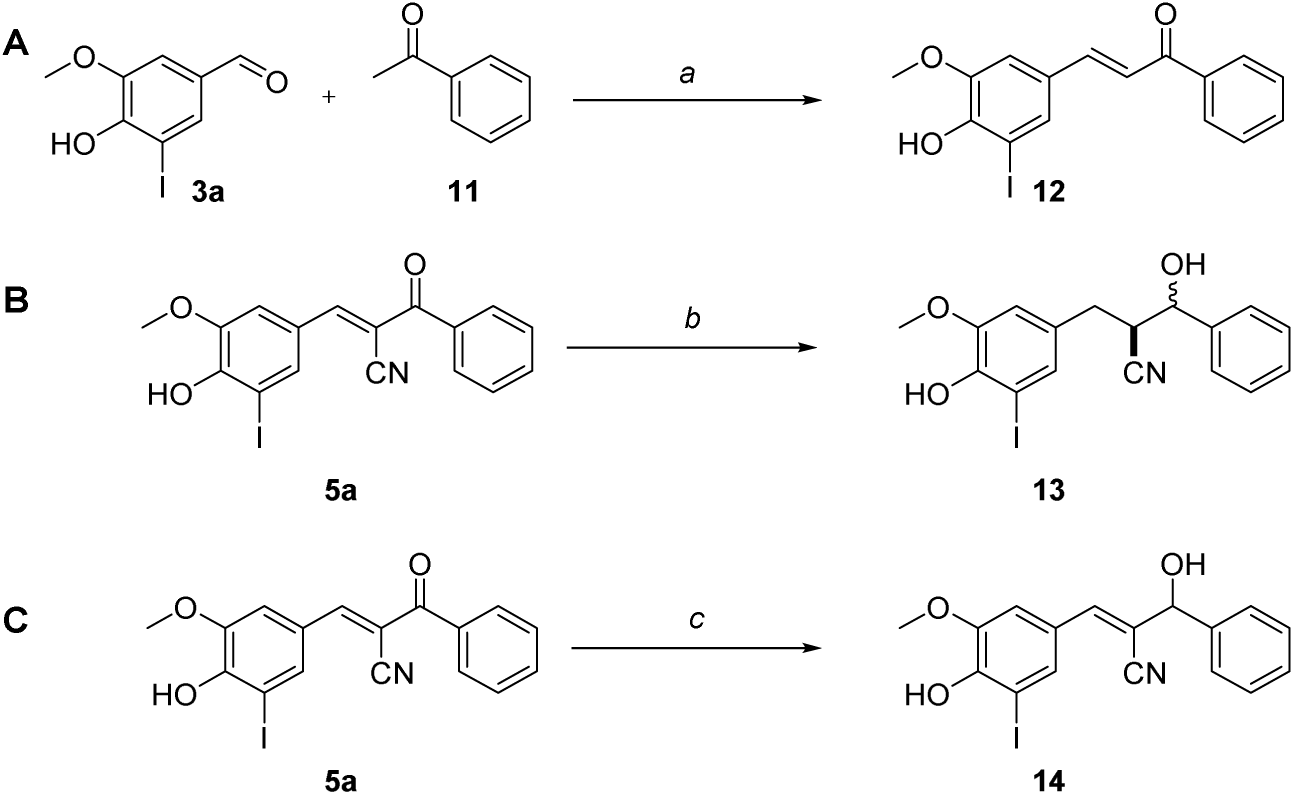
Modifications of the Michael acceptor.^*a*^. ^*a*^Reagents and conditions: (a) NaOH, EtOH, room temperature, 16h, (44–69%); (b) NaBH_4_, MeOH, 0 °C, 60 min (22%, 2:1 mixture of diastereomers); (c) NaBH_4_, CeCl_3_, MeOH, 0 °C, 1h (50%).

### SAR study

Our first aim was to see whether the catechol ring coulb be removed in order to exclude a redox mechanism for inhibition. The first series of compounds thus focused on variation of R^1^ where R^2^ was fixed as a phenyl (entries 1–14, Table 1). Compound **5a** lacking the catechol group as the R^2^ phenyl ring compared to **1** and **2**, showed a 10-fold decrease in potency (5.4 μM, entry 1) indicating that the OH-groups are not crucial for inhibition but are important for the potency. On the other hand, the parent compound (**5b**) was inactive indicating that thus the R^1^-substituents are essential for inhibition (entry 2). Restoration of the *p*-hydroxyl (**5c**) and the *m*-methoxy (**5d**) group did not result in active compounds (entries 3 and 4), while on the other hand, 3,4-dihydroxyphenyl as R^1^ (**5e**) was moderately active (32 μM, entry 5). Exchange of one hydroxy moiety to a nitro group (**5f**, 23 μM, entry 6) or a methoxy group on position 3 had no significant effect on the IC_50_ values (**5f**, 23 μM, entry 6; **5g**, 26 μM, entry 7). *N,N*-dimethylphenyl (**5h**) as R^1^ was inactive (entry 8).

The effect of the substitution pattern of the R^1^-phenyl ring was further investigated. We kept the 3-methoxy-4-hydroxy pattern constant and varied the 5-position of R^1^ (entries 9–12). Adding a methoxy-group on the 5-position (**5i**) did not result in inhibition of Dop (entry 9), while a chloride (**5j**) resulted in an active compound with poor activity (72 μM, entry 10). Changing the 3-methoxy group to an ethoxy group and have an additional 3-methoxy on the R^2^ phenyl ring (**5k**) was beneficial since the potency increased a 2-fold to 28 μM (entry 11) compared to **5j**. Changing the chloride to a bromide with R^2^ being a phenyl again (**5l**) increased the potency further to an IC_50_ of 20 μM (entry 12) compared to **5j**.

From the results so far, we concluded that the hydroxy group on the 4-position of R^1^ is essential for inhibition, while the potency is dependent on the electronic properties of the groups around the 4-position. To test this hypothesis, we tested compound **5m** that is lacking the 4-hydroxygroup compared to **5l** and **5n** in which the R^1^ substituents are shifted one place compared to **5a**. Both compounds were inactive (entries 13 and 14), supporting our hypothesis.

We continued to investigate the role of the 3,4,5-trisubstitution on R^1^ with R^2^ being 3,4-dihydroxyphenyl, like in the original two hits (entries 13–25). This change of R^2^ should deliver more potent compounds compared to the previous series. To further investigate the role of the substitution pattern on R^1^, we started with compound **10a**, which lacks the iodine moiety compared to **1**. The presence of a group on the 5-position seemed to be crucial based on the observation that **10a** was inactive (entry 13). The presence of an electron withdrawing nitro-group (**10b**) restored potency (4.1 μM, entry 15). Subsequent removal of the methoxy group on the 3-position (**10c**) led to a significant loss of potency (96 μM, entry 15), while additional removal of the 4-hydroxygroup (**10d**) again rendered an inactive compound (entry 18). This indicates that both the 3-methoxy and 4-hydroxy are essential for potent inhibitors. We thus continued investigating the effect of different substituents on the 5-position (entries 19–24).

Interestingly, when an electron donating methoxy group was placed on the 5-position (**10e**), this compound appeared to be inactive (entry 19). Electron withdrawing groups appear to be required on the 5-position. When we substituted the 5-position with a phenyl (**10f**) the compound was moderately active (41 μM, entry 20). It was however, not beneficial for the potency to add a more electron poor 4-fluorophenyl on position 5 (**10g**, entry 21). To further investigate the electronic properties, we tested analogs containing a Cl, Br or F on the 5-position (**10h**–**10j**, entries 22–24). **10h** and **10i** were slightly less potent than the original hit **1** (1.7 and 2.7 μM respectively) while fluorine (**10j**), on the other hand, resulted in further loss of potency (13 μM, entry 22). Apparently, subtle changes in acidity of the 4-hydroxyl moiety by modification of the 5-substituent, have a great impact on the potency of the compounds. The hybrid compound (**10k**) based on both **1** and **2** that has the 3-methoxy moiety replaced for a hydroxy moiety, was the most potent compound from the synthesized analogs (0.29 μM, entry 25). In conclusion, the 4-hydroxyl group on the R^1^-phenyl ring is essential for inhibition and the potency is most likely dependent on its acidity that is regulated by the 5-substituent. The two hydroxyl groups on the R^2^-phynyl ring increase the potency, but are not essential.

Having established initial SAR, we turned our attention to modification of the core part. Compound **1** contains a Michael acceptor in the form of an α,β-unsaturated ketone functionalized with a cyano group on the 3-position. The cyano group was previously designed as a modulator for the reactivity of the Michael acceptor which as a result, reacts reversibly covalent with cysteine nucleophiles whereas α,β-unsaturated ketones react irreversibly with cysteine residues.^37^ Because the inhibition of Dop by **1** was determined as fast reversible (*vide supra*), it is therefore unlikely that **1** binds reversible covalent with a nucleophilic residue of Dop. As a result, we do not expect the Michael acceptor to be a crucial structural element for inhibition. We synthesized a series of analogous compounds in which the Michael acceptor was systematically modified to verify this hypothesis (Table 2).

**Table 2.**
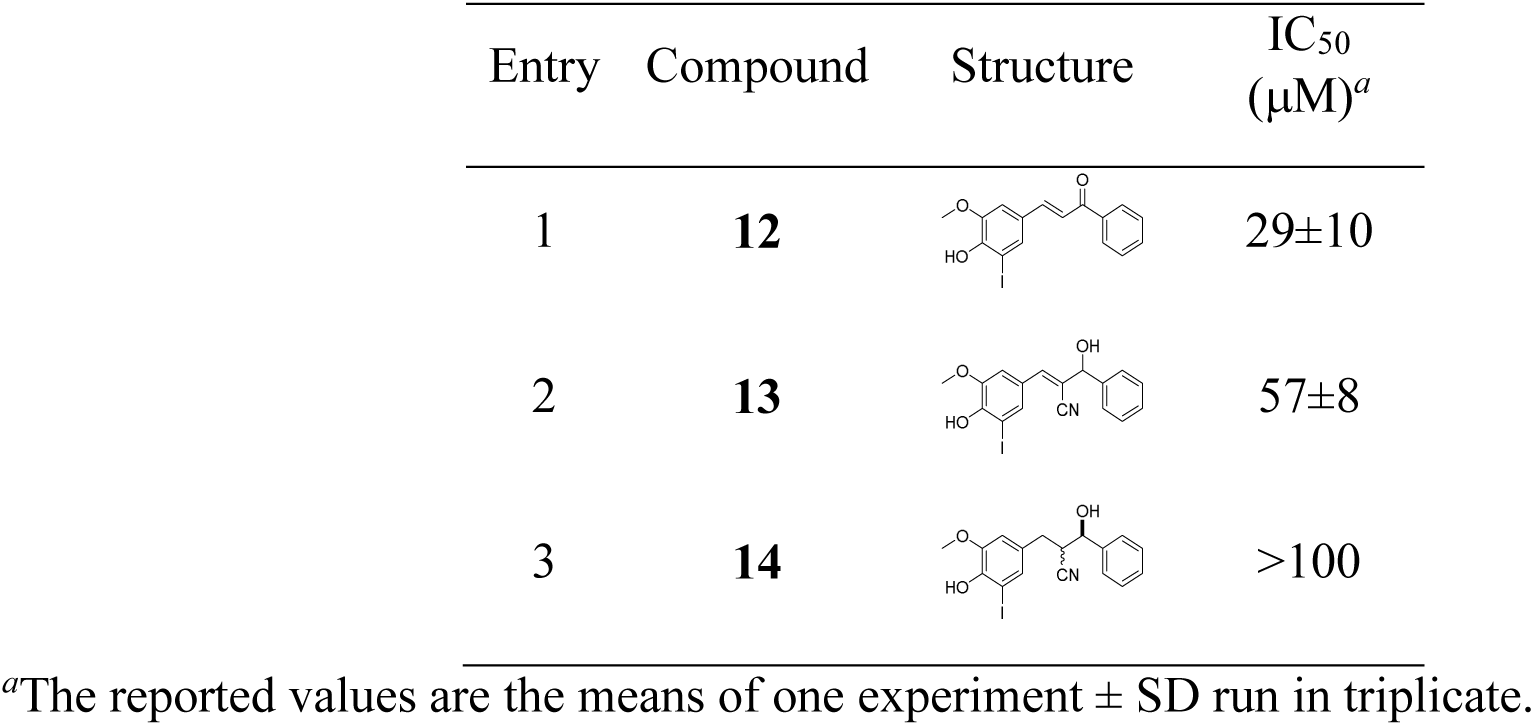
Modification of the Michael acceptor

| Entry | Compound | Structure | IC<sub>50</sub><br>(μM)<sup>a</sup> |
| --- | --- | --- | --- |
| 1 | **12** | | 29±10 |
| 2 | **13** | | 57±8 |
| 3 | **14** |  | >100 |

<sup>a</sup>The reported values are the means of one experiment ± SD run in triplicate.

Compound **12** that lacks the cyano group compared to **5a** (5.4 μM; Table 2, entry 1) was still active, but six-fold less potent (31 μM; Table 2, entry 1). On the other hand, compound **13**, in which the carbonyl is selectively reduced to the alcohol, was less potent (57 μM; entry 2). Complete reduction of the α,β-unsaturated system (**14**) resulted a complete loss of activity (entry 3). Although the full *cis*-benzylidenemalonitrile backbone is not essential for inhibition, selective reduction greatly influenced the potency of the compounds. A possible explanation could be that the rigidity of the unsaturated system is keeping the geometry of the molecule optimal for inhibition.

### Testing the compounds on native Mtb proteasome substrates

After the identification of the important structural features of the inhibitors, we tested a selection of the inhibitors on native substrates for Dop. We chose the depupylation reaction of a model pupylated substrate, malonyl coenzyme A (CoA)-acyl carrier protein transacylase (FabD∼Pup). To find the ideal conditions to test the inhibitors, we first followed the reaction over time (Figure 6B). Using 250 nM FabD∼Pup and 10 nM Dop, the depupylation reaction finished within 60 min. We chose 30 minutes as a suitable end point for the assay.

**Figure 6.**
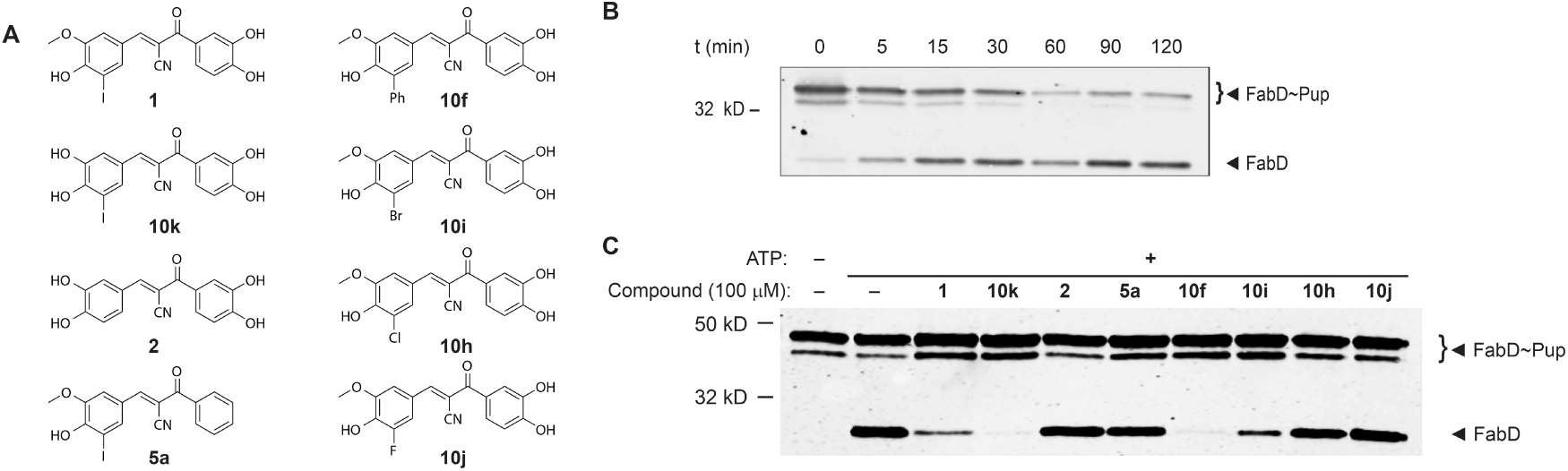
Depupylation of FabD∼Pup by Dop. A) Structures of the eight selected compounds. B) Time course of the depupylation of FabD∼Pup (250 nM) by Dop (10 nM). The samples were withdrawn at the indicated time points. B) Effect of the inhibitors (100 μM) on the depupylation reaction of FabD∼Pup (250 nM) by Dop (10 nM). The samples were withdrawn after 30 min. For B) and C), proteins were analyzed by immunoblotting using antibodies to *Mtb* FabD-His_6_.^38^ Note that FabD can be pupylated on any one of four lysines, resulting in two slightly different migration patterns.^39^

With the optimal reaction conditions determined, eight compounds were selected (Figure 6A) and tested for inhibition at 100 μM in the depupylation assay (Figure 6C). Interestingly, the degree of inhibition observed in using the fluorogenic substrate was not consistently reflected in the depupylation assays using a native substrate. For example, compound **2** (IC_50_ 0.13 µM) was identified as the most potent inhibitor, however, it failed to inhibit depupylation of FabD∼Pup by Dop. On the other hand, compound 1 (IC_50_ 0.52 µM) incompletely inhibited Dop, and **10k** (IC_50_ 0.29 µM) showed full inhibition. Compound **5a**, lacking the two phenolic OH-groups compared to **1**, had an IC_50_ of 5.4 μM in the fluorogenic assay, and failed to inhibit depupylation of FabD∼Pup. Remarkably, **10f** was not very potent in the fluorogenic assay (IC_50_ 42 μM), but fully inhibited the depupylation of FabD∼Pup. For the three compounds bearing a bromine (**10i**), chlorine (**10h**) or fluorine (**10j**) on the 5-position of the left ring, only **10i** partially inhibited Dop.

The compounds show a different inhibition pattern between the two assay substrates which are most likely explained by a difference in affinity of both substrates for Dop. It appears that the size of the 5-position of the left ring is a more important factor than the electron negativity of this substituent in the depupylation of FabD∼Pup. This is illustrated by the completely opposite effects of **2** and **10f** between the fluorogenic and native substrate (H *vs*. Ph). Moreover, the lack of inhibition of **5a** underlines the importance of the presence hydroxy groups on the right ring.

Because Dop and PafA have highly similar sequences and structures,^40^ we investigated whether or not these compounds were able to influence the ligase and depupylase activities of PafA using our pupylation substrate FabD. We looked at the pupylation of FabD by PafA in the presence of 100 μM of each of the inhibitors (Figure 7). Compounds **10k** and **2** and **10f** completely inhibited pupylation, while **1** and **10i** partly inhibited the ligase activity of PafA. The remainder of the compounds showed no inhibition. Given the structural homology, it is not surprising that inhibitors for Dop also inhibit PafA. However, **2** did not inhibit Dop in the depupylation of FabD∼Pup, while it fully inhibits PafA’s ligase activity. Apparently, despite the structural homology, small differences in inhibitor structure allow inhibitors to be selective for PafA.

**Figure 7.**
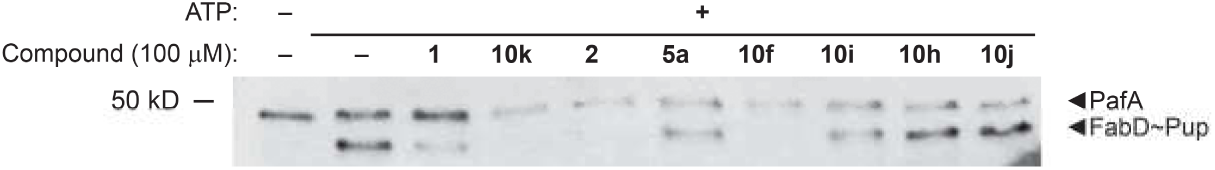
Pupylation of FabD by PafA. Effect of the inhibitors (100 μM) on the pupylation of FabD (250 nM) by PafA (50 nM). Reactions were stopped after six hours. Proteins were visualized with antibodies to FabD-His_6_, which can also detect histidine-tagged PafA used in the reactions. MW standards are indicated on the left of the blot.

Recently, we described a depupylase activity for PafA. Interestingly, PafA cannot deamidate PupQ to PupE or release AMC from Pup_(33–63)_-Glu(AMC) and appears to be more effective at depupylating inositol 1-phosphate synthetase (Ino1∼Pup) than FabD∼Pup.^21^ This observation suggests PafA and Dop may have different affinities for various native substrates. We therefore tested if the compounds could also inhibit the depupylation of either substrate by PafA (Figure 8). The same compounds that inhibited pupylation of FabD by PafA, also inhibited the depupylation of FabD and Ino1 by PafA. Compounds **10k** and **2** and **10f** completely inhibited depupylation of both Ino1∼Pup and FabD∼Pup, while **1** and **10i** partly inhibited the depupylation by PafA at 100 μM.

**Figure 8.**
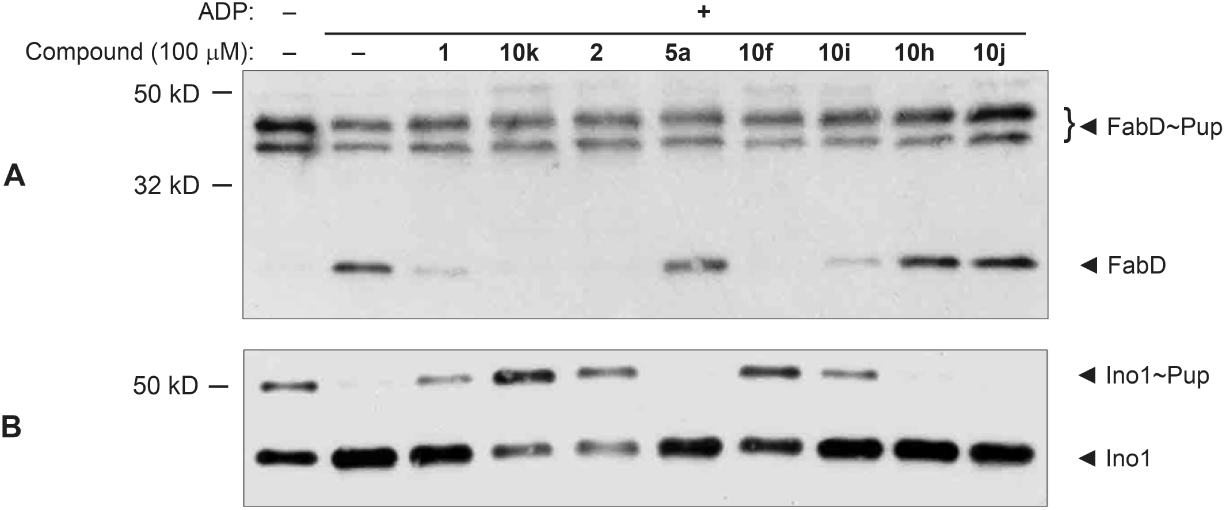
Five compounds at 100 μM inhibit depupylation of *Mtb* proteasome substrates by PafA. A) Effect of the inhibitors on the depupylation of FabD∼Pup by PafA. B) Effect of the inhibitors (100 μM) on the depupylation of Ino1∼Pup (250 nM) by PafA (50 nM). Note that Ino1 forms tetramers in which two molecules in each tetramer are pupylated; therefore, Ino1 and Ino1∼Pup co-purify as a 2:2 complex.^24^ MW standards are indicated on the left of each blot.

Both Dop and PafA are required for pupylation in *Mtb*; Dop must first deamidate PupQ to PupE before PafA ligates Pup to a substrate. The deletion of either gene product from *Mtb* results in the disappearance of the pupylome.^16^ None of the compounds was able to inhibit pupylation in *Mtb* cultures (data not shown), most likely due to the polarity of the compounds; *Mtb* has a highly hydrophobic cell envelope that is impermeable to many molecules, which contributes to the difficulty in treating TB infections.^41^ Nonetheless, our goals were to determine whether or not Dop could be targeted for inhibitor discovery as well as identify molecules that may give new insight into the mechanism of catalysis by this highly unique protease. Since our screen identified inhibitors of pupylation as well as depupylation and do not compete with ATP, these molecules may instead affect the positioning of the C-terminus of Pup and its substrate within the active site. The fact that inhibitors for both enzymes in the PPS could be found using this screening approach is likely due to the close structural homology between Dop and PafA.

A recent study showed that a well-characterized serine protease inhibitor, (4-(2-aminoethyl) benzenesulfonyl fluoride (AEBSF), could inhibit the activities of PafA.^26^ However, this molecule was not tested on Dop. This compound modified a serine in PafA that was distant from the active site. Importantly, only specific (bulky) amino acid substitutions of that residue recapitulated the effects of the inhibitor, suggesting a steric effect on Pup binding to PafA. It is possible that the inhibitors identified in our study could similarly prevent Pup binding with either Dop or PafA. Future work determining where the compounds identified in our study bind to Dop, PafA or Pup itself will almost certainly provide mechanistic insight into how these enzymes catalyze essential reactions of the PPS. Importantly our study provides the first proof-of-principle experiment that HTS can be used to identify lead compounds for Dop and PafA inhibition. Ultimately our goal will be to screen for inhibitors that will be able to better penetrate the *Mtb* cell envelope and have minimal to no toxicity in mammalian cells. Idealy, potential lead series inhibit both Dop and PafA synergistically, which could minimize the acquisition of resistance to these compounds.

## Conclusions

In an effort to identify inhibitors of Dop, we used our previously developed fluorogenic Dop substrate to successfully screen a small 1024-compound library from which compounds **1** and **2** were identified as hits. **1** was validated as a fast reversible and non-ATP competitive inhibitor for Dop and was used as starting point for a SAR-study that revealed the important structural features of the scaffold. Furthermore, we showed that several compounds were able to inhibit depupylation of FabD∼Pup by Dop, as well as pupylation and depupylation of FabD and Ino1 by PafA. Althought the inhibitors were unable to inhibit pupylation in *Mtb* cultures, we have shown that HTS is a suitable approach to identify lead compounds against *Mtb*.

## EXPERIMENTAL SECTION

### Chemistry

*General Information.* All commercially available reagents and solvents were used as purchased unless stated otherwise. Nuclear magnetic resonance (NMR) spectra were recorded on a Bruker Avance 300 (75.00 MHz for ^13^C) using the residual solvent as internal standard (^1^H: δ 7.26 ppm for CDCl_3_; 2.50 ppm for DMSO and 3.31 ppm for CD_3_OD. ^13^C{^2^H}: δ 77.00 ppm for CDCl_3_; 39.43 ppm for DMSO and 49.05 ppm for CD_3_OD). Chemical shifts (δ) are given in ppm and coupling constants (*J*) are quoted in hertz (Hz). Resonances are described as s (singlet), d (doublet), t (triplet), q (quartet), b (broad) and m (multiplet) or combinations thereof. Electrospray Ionization (ESI) high-resolution mass spectrometry was carried out using a Waters Xevo G2 XS QTOF instrument in positive ion mode (capillary potential of 3000 V) in combination with a Waters Acquity UPLC system equipped with a Waters UPLC BEH C_18_ 1.7 μm (2.1 mm x 50 mm) column using water acetonitrile mixtures.

Thin Layer Chromatography (TLC) was performed using TLC plates from Merck (SiO_2_, Kieselgel 60 F254 neutral, on aluminum with fluorescence indicator) and compounds were visualized by UV detection (254 nm) unless mentioned otherwise. All reported yields are not optimized. Flash chromatography purifications were manually performed using Grace Davisil Silica Gel (particle size 40–63 μm, pore diameter 60 Å) and the indicated eluent.

Reversed phase preparative HPLC/MS was carried out on a Waters AutoPurification system equipped with a Waters 2998 photodiode array detector, Waters 3100 mass detector and a Waters 2767 sample manager using preparative Waters X-bridge C_18_ 5 μm (30mm x 150mm or 19mm x 150mm) column in combination with water acetonitrile mixtures containing 0.1% TFA. Fractions containing the product were automatically collected based on observed mass and UV-signal after which they were lyophilized to obtain the pure products. The purity of the target compounds was determined by ^1^H-NMR analysis, high resolution mass spectroscopy and shown to be >95% pure prior to biological testing.

Compound **5k** was commercially obtained from Enamine ltd. and was declared to be >95% pure. Compound **1** was commercially obtained from Sigma Aldrich was declared to be >95% pure.

#### 3,4-bis(methoxymethoxy)benzaldehyde (**3e**)

To a stirred solution of 3,4-dihydroxybenzaldehyde (1.0g, 7.2 mmol) in acetone (30 mL) was added K_2_CO_3_ (5.0 g, 36.2 mmol) and MOMCl (1.65 mL, 21.7 mmol) at 0 °C. The reaction mixture was allowed to warm to room temperature and heated to reflux. After completion of the reaction (3 h) the suspension was filtered and the residue washed with EtOAc (25 mL). After evaporation of the solvents, the crude was purified by column chromatography (EtOAc:heptane / 1:1, R_*f*_ = 0.5) to afford **3e** (0.81g, 50%) as a white solid. ^1^H NMR (300 MHz, CDCl_3_) δ (ppm) = 9.85 (s, 1H), 7.66 (d, *J* = 1.9Hz, 1H), 7.50 (dd, *J* = 8.4, 1.9 Hz, 1H), 7.27 (d, *J* = 8.4 Hz, 1H), 5.31 (s, 2H), 5.28 (s, 2H), 3.51 (s, 3H), 3.50 (s, 3H). ^13^C NMR (75 MHz, CDCl_3_) δ (ppm) = 190.8 (CH), 152.5 (Cq), 147.4 (Cq), 131.0 (Cq), 126.3 (CH), 115.8 (CH), 115.3 (CH), 95.3 (CH_2_), 94.9 (CH_2_), 56.4 (CH_3_), 56.3 (CH_3_).

#### 5-(4-fluorophenyl)vanillin (**3p**)

To a stirred solution of 5-iodovanillin (278 mg, 1.0 mmol) and 4-fluorophenylboronic acid (284 mg, 1.5 mmol) in glycerol (10 mL), K_2_CO_3_ (152 mg, 1.1 mmol) and Pd(OAc)_2_ 22.4 mg, 0.1 mmol, 10 mol%) were added and the reaction mixture was heated in the microwave (140 °C, 20 min). The reaction mixture was added to water (150 mL) and extracted three times with EtOAc (3×50 mL). The organic layers were combined, washed with water, brine and dried over Na_2_SO_4_. Column chromatography (EtOAc:heptane / 1:1, R_*f*_ = 0.45) afforded aldehyde **3p** (194 mg, 65%) as a white solid. ^1^H NMR (300 MHz, CDCl_3_) δ (ppm) = 9.85 (s, 1H), 7.65–7.53 (m, 2H), 7.47 (d, *J* = 1.8 Hz, 1H), 7.40 (d, *J* = 1.8 Hz, 1H), 7.17–7.06 (m, 2H), 6.67 (s, 1H), 3.98 (s, 3H). ^13^C NMR (75 MHz, CDCl_3_) δ (ppm) = 190.9 (CH), 162.2 (d, *J*C–F = 147 Hz, CF) 148.6 (Cq), 147.4 (Cq), 132.2 (Cq), 130.7 (CH), 130.6 (CH), 129.1 (Cq), 128.2 (CH), 126.5 (Cq), 115.3 (CH), 115.1 (CH), 107.5 (CH), 56.3 (CH_3_). ^19^F NMR (282 MHz, CDCl_3_) δ (ppm) = –114.3.

#### 3-(3,4-bis(methoxymethoxy)phenyl)-3-oxopropanenitrile (**8**)

Acetonitrile (1.63 mL, 31 mmol) was dissolved in dry THF (40 mL) and the solution was cooled to –78 °C. *n*-Buli (19.5 mL, 1.6M in hexanes) was added dropwise *via* a syringe. The white suspension was stirred 1h at this temperature after which a solution of 3,4-bis(methoxymethoxy)benzoic acid methyl ester (2.0g 7.8 mmol) in THF (5 mL) was added drop wise. After stirring for 1h at –78 °C, hythe reaction was quenched by adding sat. NH_4_Cl (10 mL) and the mixture allowed to warm to room temperature and water (50 mL) was added. The aqueous layer was extracted three times with EtOAc (30 mL). The organic layers were combined, washed with brine, dried over Na_2_SO_4_ and concentrated *in vacuo* to give **8** (1.96g, 95%) as a white solid. ^1^H NMR (300 MHz, CDCl_3_) δ (ppm) = 7.75 (d, *J* = 2.1 Hz, 1H), 7.55 (dd, *J* = 8.6, 2.2 Hz, 1H), 7.24 (d, *J* = 8.6 Hz, 1H), 5.32 (s, 2H), 5.29 (3, 2H), 4.01 (s, 2H), 3.53 (s, 3H), 3.52 (s, 3H). ^13^C NMR (75 MHz, CDCl_3_) δ (ppm) = 185.4 (Cq), 152.8 (Cq), 147.2 (Cq), 128.5 (Cq), 124.1 (CH), 116.2 (CH), 115.1 (CH), 113.9 (Cq), 95.4 (CH_2_), 94.9 (CH_2_), 56.6 (CH_3_), 56.5 (CH_3_), 29.1 (CH_2_).

#### 5-phenylvanillin (**3o**)

To a stirred solution of 5-iodovanillin (278 mg, 1.0 mmol) and 4-phenylboronic acid (182 mg, 1.5 mmol) in glycerol (10 mL), K_2_CO_3_ (152 mg, 1.1 mmol) and Pd(OAc)_2_ (22.4 mg, 0.1 mmol, 10 mol%) were added and the reaction mixture was heated in the microwave (140 °C, 20 min). The reaction mixture was added to water (150 mL) and extracted three times with EtOAc (3×50 mL). The organic layers were combined, washed with water, brine and dried over Na_2_SO_4_. Column chromatography (EtOAc:heptane / 1:1, R_*f*_ = 0.45) afforded aldehyde **3o** (151 mg, 66%) as a white solid. ^1^H NMR (300 MHz, CDCl_3_) δ (ppm) = 9.87 (s, 1H), 7.65–7.56 (m, 2H), 7.49 (d, *J* = 1.8 Hz, 1H), 7.42 (d, *J* = 1.8 Hz, 1H), 7.20–7.09 (m, 2H), 6.48 (s, 1H), 4.02 (s, 3H) ^13^C NMR (75 MHz, CDCl_3_) δ (ppm) = 191.0 (CH), 148.6 (Cq), 147.4 (Cq), 136.2 (Cq), 129.1 (Cq), 129.0 (2xCH), 128.6 (CH), 128.3 (2xCH), 127.7 (CH), 127.6 (Cq), 107.426.3 (CH_3_).

### General procedure A, Knoevenagel condensation

The appropriate 2-oxoacetonitrile (1 equiv) and aldehyde (1 equiv) were dissolved in absolute EtOH. A catalytic amount of piperidine was added and the mixture was heated to 65 °C for 30 minutes. The mixture was cooled to room temperature and stirred overnight. The precipitated product was then filtered and the filtrate washed with ice-cold EtOH to obtain the crude product.

### General procedure B, MOM-deprotection

The MOM-protected compound (1 equiv) was dissolved in CH_2_Cl_2_ followed by the addition of a scoop of KHSO_4_ supported silicagel.^36^ The resulting suspension was stirred overnight and the silica was filtered off. The residu was evaporated and the crude purified using preparative LCMS.

### General Procedure C, Chalchone formation

To a stirred solution of KOH (45 equiv) in water cooled to 0 °C, was added dropwise a solution of benzophenone (1.5 equiv) and benzaldehyde (1.0 equiv) in MeOH. The mixture was kept at 0 °C for 3 hours after which the mixture was stirred overnight at room temperature. The mixture was poured in ice and the pH was adjusted to 3–4 with 1M HCl and extracted three times with EtOAc. The organic layers were combined, washed with brine, dried over MgSO_4_ and evaporated to obtain the crude product.

#### (E)-2-benzoyl-3-(4-hydroxy-3-iodo-5-methoxyphenyl)acrylonitrile (**5a**)

According to general procedure A, the reaction between benzoylacetonitrile (0.78 g, 5.4 mmol) and 5-idoovanillin (1.5 g, 5.4 mmol) afforded product **5a** (1.87 g, 85%) as an orange solid. ^1^H NMR (300 MHz, DMSO) δ (ppm) = 11.01 (bs, 1H), 8.15 (d, *J* = 1.9 Hz, 1H), 8.01 (s, 1H), 7.85–7.77 (m, 3H), 7.74–7.65 (m, 1H), 7.63–7.53 (m, 2H), 3.87(s, 3H). ^13^C NMR (75 MHz, DMSO) δ (ppm) = 189.9 (Cq), 154.3 (CH), 151.5 (Cq), 146.7 (Cq), 136.0 (Cq), 135.2 (CH), 132.9 (CH), 129.1 (2xCH), 128.6 (2xCH), 124.8 (Cq), 117.3 (Cq), 106.8 (CH), 84.9 (Cq), 56.1 (CH_3_). HRMS ESI (3000V): calculated for C_17_H_13_NO_3_I (M+H^+^) 405.9940, found 405.9948.

#### (E)-2-benzoyl-3-phenylacrylonitrile (**5b**)

According to general procedure A, the reaction between benzoylacetonitrile (1.45 g, 10 mmol) and benzaldehyde (1.06 g, 10 mmol) afforded product **5b** (2.04g, 87%) as an off-white solid. ^1^H NMR (300 MHz, CDCl_3_) δ (ppm) = 8.07 (s, 1H), 8.05–8.00 (m, 2H), 7.96–7.86 (m, 2H), 7.69–7.48 (m, 6H). ^13^C NMR (75 MHz, CDCl_3_) δ (ppm) = 188.9 (Cq), 155.5 (CH), 135.7 (Cq), 133.4 (Cq), 133.3 (Cq), 131.8 (Cq), 131.1 (2xCH), 129.3 (2xCq), 129.2 (2xCH), 128.8 (2xCH), 116.8 (Cq), 110.1 (Cq). HRMS ESI (3000V): calculated for C_16_H_12_NO (M+H^+^) 234.0913, found 234.0907.

#### (E)-2-benzoyl-3-(4-hydroxyphenyl)acrylonitrile (**5c**)

According to general procedure A, the reaction between benzoylacetonitrile (0.145 g, 1.0 mmol) and 4-hydroxybenzaldehyde (0.106 g, 1.0 mmol) afforded product **5c** (0.189g, 76%) as a yellow solid. ^1^H NMR (300 MHz, DMSO) δ (ppm) = 8.08–7.96 (m, 3H), 7.84–7.76 (m, 2H), 7.73–7.63 (m, 1H), 7.72–7.51 (m, 2H), 7.02– 6.90 (m, 2H). ^13^C NMR (75 MHz, DMSO) δ (ppm) = 119.7 (Cq), 172.7 (Cq), 165.2 (CH), 145.8 (Cq), 143.7 (2xCH), 142.2 (CH), 138.6 (2xCH), 138.2 (2xCH), 132.3 (Cq), 127.2 (Cq), 126.0 (2xCH), 114.8 (Cq). HRMS ESI (3000V): calculated for C_16_H_12_NO_2_ (M+H^+^) 250.0868, found 250.0873.

#### (E)-2-benzoyl-3-(4-hydroxy-3-methoxyphenyl)acrylonitrile (**5d**)

According to general procedure A, the reaction between benzoylacetonitrile (0.145 g, 1.0 mmol) and 3-methoxy-4-hydroxybenzaldehyde (0.122 g, 1.0 mmol) afforded product **5d** (0.233g, 83%) as a yellow solid. ^1^H NMR (300 MHz, CDCl3) δ (ppm) = 7.98 (s, 1H), 7.91 (d, *J* = 2.1 Hz, 1H), 7.88–7.81 (m, 2H), 7.66–7.56 (m, 1H), 7.55–7.46 (m, 1H), 7.39 (dd, *J* = 8.6, 2.1 Hz, 1H), 6.95 (d, *J* = 8.3 Hz, 1H), 5.12 (bs, 1H), 3.96 (s, 3H). ^13^C NMR (75 MHz, CDCl_3_) δ (ppm) = 189.4 (Cq), 155.7 (CH), 151.1 (Cq), 146.9 (Cq), 136.3 (Cq), 133.0 (CH), 129.1 (2xCH), 128.6 (2xCH), 124.6 (Cq), 117.9 (Cq), 115.0 (CH), 111.2 (CH), 106.4 (Cq), 56.2 (CH_3_). HRMS ESI (3000V): calculated for C_17_H_14_NO_3_ (M+H^+^) 280.0968, found 280.0974.

#### (E)-2-benzoyl-3-(3,4-dihydroxyphenyl)acrylonitrile (**5e**)

According to general procedure A, the reaction between benzoylacetonitrile (0.10 g, 0.44 mmol) and *bis*-MOM-protected 3,4-dihydroxybenzaldehyde (0.071 g, 0.49 mmol) afforded the crude product (121 mg) of which 20 mg was directly used in the deprotection according to general procedure B. This afforded **5e** (13 mg, 87%) as an orange solid.^1^H NMR (300 MHz, CD_3_OD) δ (ppm) = 7.82 (s, 1H), 7.73– 7.66 (m, 2H), 7.62 (d, *J* = 2.3 Hz, 1H), 7.58–7.51 (m, 1H), 7.48–7.40 (m, 2H), 7.29 (ddd, *J* = 8.4, 2.3, 0.3 Hz, 1H), 6.80 (d, *J* = 8.3 Hz, 1H). ^13^C NMR (75 MHz, CD_3_OD) δ (ppm) = 190.8 (Cq), 171.6 (Cq), 156.3 (CH), 152.0 (Cq), 145.8 (Cq), 136.7 (Cq), 132.5 (CH), 128.8 (2xCH), 128.3 (2xCH), 127.0 (CH), 123.9 (Cq), 117.1 (Cq), 116.4 (CH), 115.4 (CH), 105.3 (Cq). HRMS ESI (3000V): calculated for C_16_H_12_NO_3_ (M+H^+^) 266.0812, found 266.0815.

#### (E)-2-benzoyl-3-(4-hydroxy-3-methoxy-5-nitrophenyl)acrylonitrile (**5f**)

According to general procedure A, the reaction between benzoylacetonitrile (73 mg, 0.5 mmol) and 5-nitrovanilin (99 mg, 0.5 mmol) afforded product **5f** (139mg, 94%) as a yellow solid. ^1^H NMR (300 MHz, CDCl_3_) δ (ppm) = 8.08 (d, *J* = 2.2 Hz, 1H), 8.01 (d, *J* = 2.3Hz, 1H), 7.91 (s, 1H), 7.86–7.78 (m, 2H), 7.66–7.56 (m, 1H), 7.56–7.46 (m, 2H), 3.94 (s, 3H). ^13^C NMR (75 MHz, CDCl_3_) δ (ppm) = 190.0 (Cq), 161.4 (Cq), 155.2 (CH), 153.9 (Cq), 136.8 (Cq), 136.3 (Cq), 132.7 (CH), 129.1 (CH), 129.0 (2xCH), 128.5 (2xCH), 118.5 (Cq), 115.9 (Cq), 110.4 (CH), 103.9 (Cq), 56.4 (CH_3_). HRMS ESI (3000V): calculated for C_17_H_13_N_2_O_5_ (M+H^+^) 325.0825, found 325.0814.

#### (E)-2-benzoyl-3-(4-hydroxy-3-nitrophenyl)acrylonitrile (**5g**)

According to general procedure A, the reaction between benzoylacetonitrile (290 mg, 2.0 mmol) and 4-hydroxy-3-nitrobenzaldehyde (334 mg, 2.0 mmol) afforded **5g** (55 mg, 92%) as an pale yellow solid. ^1^H NMR (300 MHz, DMSO) δ (ppm) = 8.67 (d, *J* = 2.3 Hz, 1H), 8.28 (dd, *J* = 8.9, 2.3 Hz, 1H), 8.14 (s, 1H), 7.88–7.80 (m, 2H), 7.75–7.66 (m, 1H), 7.63–7.52 (m, 2H), 7.31 (d, *J* = 8.8 Hz, 1H). ^13^C NMR (75 MHz, DMSO) δ (ppm) = 189.5 (Cq), 155.8 (Cq), 153.4 (CH), 137.1 (Cq), 136.3 (CH), 135.6 (Cq), 133.1 (CH), 129.2 (2xCH), 128.6 (2xCH), 122.7 (Cq), 119.9 (CH), 116.47 (Cq), 108.7 (Cq). HRMS ESI (3000V): calculated for C_16_H_11_N_2_O_4_ (M+H^+^) 295.0713, found 295.0702.

#### (E)-2-benzoyl-3-(3-(dimethylamino)phenyl)acrylonitrile (**5h**)

According to general procedure A, the reaction between benzoylacetonitrile (290 mg, 2.0 mmol) and 4-dimethylaminobenzaldehyde (298 mg, 2.0 mmol) afforded after filtration **5h** (420 mg, 76%) as an bright orange solid. ^1^H NMR (300 MHz, DMSO) δ (ppm) = 8.05–7.96 (m, 2H), 7.50 (s, 1H), 7.78–7.72 (m, 2H), 7.69–7.60 (m, 1H), 7.59–7.50 (m, 1H), 6.91–6.80 (m, 2H), 3.10 (s, 6H). ^13^C NMR (75 MHz, DMSO) δ (ppm) = 190.1 (Cq), 155.3 (CH), 153.9 (Cq), 137.1 (Cq), 134.1 (2xCH), 132.2 (CH), 128.6 (2xCH), 128.4 (2xCH), 118.8 (Cq), 118.5 (Cq), 111.7 (2xCH), 100.4 Cq), 39.6 (2xCH3). HRMS ESI (3000V): calculated for C_18_H_17_NO_2_ (M+H^+^) 277.1335, found 277.1333.

#### (E)-2-benzoyl-3-(4-hydroxy-3,5-dimethoxyphenyl)acrylonitrile (**5i**)

According to general procedure A, the reaction between benzoylacetonitrile (0.145 g, 1.0 mmol) and 3,5-dimethoxy-4-hydroxybenzaldehyde (0.182 g, 1.0 mmol) afforded product **5i** (260 mg, 84%) as a yellow solid. ^1^H NMR (300 MHz, CDCl_3_) δ (ppm) = 7.99 (s, 1H), 7.92–7.85 (m, 2H), 7.67–7.58 (m, 1H), 7.56–7.48 (m, 2H), 7.40 (s, 2H), 3.97 (s, 6H). ^13^C NMR (75 MHz, CDCl_3_) δ (ppm) = 189.3 (Cq), 155.8 (CH), 147.2 (Cq), 140.3 (Cq), 136.2 (Cq), 133.1 (CH), 129.1 (2xCH), 128.6 (2xCH), 123.3 (Cq), 117.8 (Cq), 108.7 (2xCH), 106.7 (Cq), 56.6 (2xCH_3_). HRMS ESI (3000V): calculated for C_18_H_16_NO_4_ (M+H^+^) 310.1074, found 310.1085.

#### (E)-2-benzoyl-3-(3-chloro-4-hydroxy-5-methoxyphenyl)acrylonitrile (**5j**)

According to general procedure A, the reaction between benzoylacetonitrile (73 mg, 0.5 mmol) and 5-chlorovanillin (93 mg, 0.5 mmol) afforded product **5j** (125 mg, 80%) as a yellow solid. ^1^H NMR (300 MHz, CDCl_3_) δ (ppm) = 7.91 (s, 1H), 7.89 (d, *J* = 2.2 Hz, 1H), 7.87–7.80 (m, 2H), 7.65–7.56 (m, 1H), 7.54–7.43 (m, 3H), 3.95 (s, 3H). ^13^C NMR (75 MHz, CDCl_3_) δ (ppm) = 189.7 (Cq), 154.8 (CH), 153.8 (Cq), 149.4 (Cq), 136.7 (Cq), 132.8 (CH), 130.0 (CH), 129.1 (2xCH), 128.6 (2xCH), 121.3 (Cq), 120.8 (Cq), 118.4 (Cq), 109.5 (CH), 104.5 (Cq), 56.4 (CH_3_). HRMS ESI (3000V): calculated for C_17_H_13_NO_3_Cl (M+H^+^) 314.0584, found 314.0582.

#### (E)-2-benzoyl-3-(3-bromo-4-hydroxy-5-methoxyphenyl)acrylonitrile (**5i**)

According to general procedure A, the reaction between benzoylacetonitrile (73 mg, 0.5 mmol) and 5-bromovanillin (116 mg, 0.5 mmol) afforded product **5i** (64 mg, 36%) as an orange solid. ^1^H NMR (300 MHz, CD_3_OD) δ (ppm) = 7.95 (s, 1H), 7.85–7.78 (m, 3H), 7.77–7.75 (m, 1H), 7.69–7.61 (m, 1H), 7.60–7.50 (m, 2H), 3.94 (s, 3H). ^13^C NMR (75 MHz, CD_3_OD) δ (ppm) = 190.3 (Cq), 154.7 (CH), 148.6 (Cq), 136.4 (Cq), 132.7 (CH), 130.9 (CH), 128.9 (2xCH), 128.4 (2xCH), 123.4 (Cq), 117.2 (Cq), 111.1 (CH), 109.7 (Cq), 106.2 (Cq), 55.5 (CH_3_). HRMS ESI (3000V): calculated for C_17_H_13_NO_3_Br (M+H^+^) 358.0079, found 358.0093.

#### (E)-2-benzoyl-3-(3-bromo-5-methoxyphenyl)acrylonitrile (**5m**)

According to general procedure A, the reaction between benzoylacetonitrile (73 mg, 0.5 mmol) and 3-bromo-5-methoxybenzaldehyde (108 mg, 0.5 mmol) afforded product **5m** (73 mg, 43%) as a yellow solid. ^1^H NMR (300 MHz, CDCl_3_) δ (ppm) = 7.97–7.876 (m, 3H), 7.72–7.61 (m, 2H), 7.72– 7.48 (m, 3H), 7.29–7.23 (m, 1H), 3.88 (s, 3H). ^13^C NMR (75 MHz, CDCl_3_) δ (ppm) = 188.4 (Cq), 160.7 (Cq), 153.6 (CH), 135.4 (Cq), 134.2 (Cq), 129.4 (2xCH), 128.8 (2xCH), 126.7 (CH), 123.5 (Cq), 122.5 (CH), 113.6 (Cq), 113.4 (CH), 111.7 (Cq), 55.8 (CH_3_). HRMS ESI (3000V): calculated for C_17_H_13_NO_2_Br (M+Na^+^) 363.9949, found 363.9975.

#### (E)-2-benzoyl-3-(3-hydroxy-2-iodo-4-methoxyphenyl)acrylonitrile (**5n**)

According to general procedure A, the reaction between benzoylacetonitrile (146 mg, 1 mmol) and 2-iodo-3-hydroxy-4-methoxybenzaldehyde (278 mg, 1 mmol) afforded product **5n** (351 mg, 87%) as a yellow solid. ^1^H NMR (300 MHz, DMSO) δ (ppm) = 10.08 (bs, 1H), 8.19 (s, 1H), 7.92–7.79 (m, 3H), 7.76–7.66 (m, 1H), 7.65–7.54 (m, 2H), 7.25 (d, *J* = 8.7 Hz, 2H), 3.94 (s, 3H). ^13^C NMR (75 MHz, DMSO) δ (ppm) = 189.8 (Cq), 159.2 (CH), 150.4 (Cq), 147.0 (Cq), 135.7 (Cq), 129.2 (2xCH), 128.7 (2xCH), 127.2 (Cq), 122.0 (CH), 116.2 (Cq), 111.3 (CH), 110.2 (Cq), 93.9 (Cq), 56.4 (CH_3_). HRMS ESI (3000V): calculated for C_17_H_13_NO_3_I (M+H^+^) 405.9940, found 405.9948.

#### (E)-2-(3,4-dihydroxybenzoyl)-3-(4-hydroxy-3-methoxyphenyl)acrylonitrile (**10a**)

According to general procedure A, the reaction between malonitrile **8** (26.5 mg, 0.10 mmol) and vanillin (15.2 mg, 0.10 mmol) afforded the crude product that was used without purification in the deprotection according to general procedure B. This afforded **10a** (13 mg, 41%) as an orange solid. ^1^H NMR (300 MHz, CD_3_OD) δ (ppm) = 7.95 (s, 1H), 7.86 (d, *J* = 2.1 Hz, 1H), 7.48 (dd, *J* = 8.6, 2.1 Hz, 1H), 7.35–7.28 (m, 3H), 6.92 (d, *J* = 8.3 Hz, 1H), 6.91–6.85 (m, 1H), 3.93 (s, 3H). ^13^C NMR (75 MHz, CD_3_OD) δ (ppm) = 190.6 (Cq) 156.5 (CH), 153.8 (Cq), 152.5 (Cq), 149.4 (Cq), 146.7 (Cq), 129.3 (Cq) 129.1 (CH), 125.4 (Cq), 124.4 (CH), 119.1 (Cq), 117.4 (CH), 116.9 (CH), 115.9 (CH), 113.8 (CH), 107.2 (Cq), 56.5 (CH_3_). HRMS ESI (3000V): calculated for C_17_H_14_NO_5_ (M+H^+^) 312.0872, found 312.0870.

#### (E)-2-(3,4-dihydroxybenzoyl)-3-(4-hydroxy-3-methoxy-5-nitrophenyl)acrylonitrile (**10b**)

According to general procedure A, the reaction between malonitrile **8** (26.5 mg, 0.10 mmol) and 5-nitrovanillin (19.7 mg, 0.10 mmol) afforded the crude product that was used without purification in the deprotection according to general procedure B. This afforded **10b** (21 mg, 59%) as a yellow solid. ^1^H NMR (300 MHz, CD_3_OD) δ (ppm) = 8.21 (d, *J* = 2.0 Hz, 1H), 8.02 (d, *J* = 2.0 Hz, 1H), 7.97 (s, 1H), 7.40–7.30 (m, 2H), 6.89 (d, *J* = 8.1 Hz, 1H), 3.99 (s, 3H). ^13^C NMR (75 MHz, CD_3_OD) δ (ppm) = 189.4 (Cq), 153.8 (CH), 152.9 (Cq), 151.7 (Cq), 146.8 (Cq), 137.6 (Cq), 128.7 (Cq), 124.8 (CH), 123.7 (Cq), 122.6 (CH), 118.3, (Cq), 117.4 (Cq), 117.4 (CH), 116.4 (CH), 116.0 (CH), 110.5 (Cq), 57.4 (CH_3_). HRMS ESI (3000V): calculated for C_17_H_13_N_2_O_7_ (M+H^+^) 357.0723, found 357.0731.

#### (E)-2-(3,4-dihydroxybenzoyl)-3-(4-hydroxy-3-nitrophenyl)acrylonitrile (**10c**)

According to general procedure A, the reaction between malonitrile **8** (20.0 mg, 0.075 mmol) and 4-dimethylaminobenzaldehyde (12.8 mg, 0.075 mmol) afforded the crude product that was used without purification in the deprotection according to general procedure B. This afforded **10c** (14 mg, 71%) as an orange solid. ^1^H NMR (300 MHz, CD_3_OD) δ (ppm) = 8.73 (d, *J* = 2.3 Hz, 1H), 8.25 (dd, *J* = 9.0, 2.4 Hz, 1H), 7.98 (s, 1H), 7.40–7.30 (m, 2H), 7.19 (d, *J* = 8.9 Hz, 1H). ^13^C NMR (75 MHz, CD_3_OD) δ (ppm) = 189.7 (Cq), 153.5 (CH), 152.8 (Cq), 146.8 (Cq), 138.2 (CH), 137.4 (Cq), 130.5 (CH), 128.9 (Cq), 124.7 (CH), 122.96 (CH), 118.2(Cq), 117.4 CH), 116.0 (CH). HRMS ESI (3000V): calculated for C_16_H_11_N_2_O_6_ (M+H^+^) 327.0612, found 327.0615.

#### (E)-2-(3,4-dihydroxybenzoyl)-3-(3-nitrophenyl)acrylonitrile (**10d**)

According to general procedure A, the reaction between malonitrile **8** (20.0 mg, 0.075 mmol) and 3-nitrobenzaldehyde (12.8 mg, 0.075 mmol) afforded the crude product that was used without purification in the deprotection according to general procedure B. This afforded **10d** (15 mg, 65%) as a yellow solid. ^1^H NMR (300 MHz, CD_3_OD) δ (ppm) = 8.92 (t, *J* = 2.1 Hz, 1H), 8.45 (ddd, *J* = 8.3, 3.2, 0.9 Hz, 1H), 8.41–8.36 (m, 1H), 8.13 (s, 1H), 7.83 (t, *J* = 8.1 Hz, 1H), 7.58– 7.40 (m, 2H), 6.93 (d, *J* = 8.2 Hz, 1H).^13^C NMR (75 MHz, CD_3_CN) δ (ppm) = 188.4 (Cq), 152.4 (CH), 151.5 (Cq), 149.4 (Cq), 136.9 (CH), 134.6 (Cq), 131.4 (CH), 128.4 (Cq), 125.5 (CH), 124.9 (CH), 118.2 (CH), 117.4 (Cq), 116.0 (CH), 114.7 (Cq). HRMS ESI (3000V): calculated for C_16_H_11_N_2_O_5_ (M+H^+^) 311.0662, found 311.0675.

#### (E)-2-(3,4-dihydroxybenzoyl)-3-(4-hydroxy-3,5-dimethoxyphenyl)acrylonitrile (**10e**)

According to general procedure A, the reaction between malonitrile **8** (26.5 mg, 0.10 mmol) and 5-methoxyvanillin (18.2 mg, 0.10 mmol) afforded the crude product that was used without purification in the deprotection according to general procedure B. This afforded **10e** (7 mg, 20%) as a yellow solid. ^1^H NMR (300 MHz, CD_3_OD) δ (ppm) = 7.96 (s, 1H), 7.47 (s, 1H), 7.36–7.28 (m, 3H), 6.93–6.85 (m, 1H), 3.91 (s, 6H). ^13^C NMR (75 MHz, CD_3_OD) δ (ppm) = 190.5 (Cq), 156.7 (CH), 152.5 (Cq), 149.5 (Cq), 146.7 (Cq), 129.3 (Cq), 124.5 (CH), 124.4 (Cq), 117.4 (CH), 115.9 (CH), 110.1 (2xCH), 107.6 (Cq), 56.9 (2xCH_3_). HRMS ESI (3000V): calculated for C_18_H_16_NO_6_ (M+H^+^) 342.0978, found 342.0982.

#### (E)-2-(3,4-dihydroxybenzoyl)-3-(6-hydroxy-5-methoxy-[1,1’-biphenyl]-3-yl)acrylonitrile (**10f**)

According to general procedure A, the reaction between malonitrile **8** (29.1 mg, 0.11 mmol) and 5-phenylvanillin (25.0 mg, 0.11 mmol) afforded the crude product that was used without purification in the deprotection according to general procedure B. This afforded **10f** (24 mg, 57%) as an orange solid. ^1^H NMR (300 MHz, CD_3_OD) δ (ppm) = 7.98 (s, 1H), 7.80 (d, *J* = 2.1Hz, 1H) 7.61–7.54 (m, 3H), 7.42–7.25 (m, 6H), 6.88 (d, *J* = 8.8 Hz, 1H), 3.96 (s, 3H). ^13^C NMR (75 MHz, CD_3_OD) δ (ppm) = 190.4 (Cq), 156.6 (CH), 152.5 (Cq), 150.6 (Cq), 149.6 (Cq), 146.6 (Cq), 138.5 (Cq), 130.4 (2xCH), 130.2 (Cq), 129.8 (CH), 129.2 (Cq), 129.1 (2xCH), 128.4 (CH), 124.9 (Cq), 124.5 (CH), 119.1 (Cq), 117.5 (CH), 115.9 (CH) 112.2 (CH), 107.6 Cq), 56.8 (CH_3_). HRMS ESI (3000V): calculated for C_23_H_18_NO_5_ (M+H^+^) 388.1179, found 388.1199.

#### (E)-2-(3,4-dihydroxybenzoyl)-3-(4’-fluoro-6-hydroxy-5-methoxy-[1,1’-biphenyl]-3-yl)acrylonitrile (**10g**)

According to general procedure A, the reaction between malonitrile **8** (22.4 mg, 0.08 mmol) and 5-(4-fluorophenyl)vanillin (25.0 mg, 0.08 mmol) afforded the crude product that was used without purification in the deprotection according to general procedure B. This afforded **10g** (9 mg, 23%) as an orange solid. ^1^H NMR (300 MHz, CD_3_OD) δ (ppm) = 7.99 (s, 1H), 7.82 (d, *J* = 2.1 Hz, 7.36–1H), 7.65–7.56 (m, 3H), 7.36–7.30 (m, 2H), 7.18–7.06 (m, 2H), 6.93–6.85 (m, 1H), 3.98 (s, 3H). ^13^C NMR (75 MHz, CD_3_OD) δ (ppm) = 188.9 (Cq), 163.8 (Cq), 160.6 (Cq), 155.1 (CH), 151.1 (Cq), 149.1 (Cq), 148.2 (Cq), 145.3 (Cq), 133.2 (d, *J*_*C-F*_ = 3.3 Hz, CF), 130.9 (CH), 130.8 (CH), 128.2 (CH), 127.8 (Cq), 127.6 (Cq), 123.5 (Cq), 123.1 (CH), 117.7 (Cq), 116.0 (CH), 114.6 (CH), 114.5 (CH), 114.3 (CH), 110.9 (CH), 106.3 (Cq), 55.4 (CH_3_). ^19^F NMR (282 MHz, CD_3_OD) δ (ppm) = –117.38. HRMS ESI (3000V): calculated for C_23_H_17_NO_5_F (M+H^+^) 406.1091, found 406.1080.

#### (E)-3-(3-chloro-4-hydroxy-5-methoxyphenyl)-2-(3,4-dihydroxybenzoyl)acrylonitrile (**10h**)

According to general procedure A, the reaction between malonitrile **8** (20.0 mg, 0.075 mmol) and 5-chlorovanilin (14.7 mg, 0.075 mmol) afforded the crude product that was used without purification in the deprotection according to general procedure B. This afforded **10h** (20 mg, 77%) as an orange solid. ^1^H NMR (300 MHz, CD_3_OD) δ (ppm) = 7.90 (s, 1H), 7.76 (d, *J* = 2.1 Hz, 1H), 7.65 (d, *J* = 1.8 Hz, 1H), 7.38–7.29 (m, 2H), 6.94–6.85 (m, 1H), 3.95 (s, 3H). ^13^C NMR (75 MHz, CDCl_3_) δ (ppm) = 190.0 (Cq), 154.8 (CH), 152.7 (Cq), 150.7 (Cq), 149.7 (Cq), 146.7 (Cq), 131.2 (CH), 129.0 (Cq), 125.6 (Cq), 124.6 (CH), 118.7 (Cq), 117.4 (CH), 116.0 (CH), 112.6 (CH), 110.6 (Cq), 108.7 (Cq), 57.0 (CH_3_). HRMS ESI (3000V): calculated for C_17_H_13_NO_5_Cl (M+H^+^) 346.0477, found 346.0471.

#### (E)-3-(3-bromo-4-hydroxy-5-methoxyphenyl)-2-(3,4-dihydroxybenzoyl)acrylonitrile (**10i**)

According to general procedure A, the reaction between malonitrile **8** (20.0 mg, 0.075 mmol) and 5-bromovanilin (17.4 mg, 0.075 mmol) afforded the crude product that was used without purification in the deprotection according to general procedure B. This afforded **10i** (18 mg, 61%) as an orange solid. ^1^H NMR (300 MHz, CD_3_OD) δ (ppm) = 7.88 (s, 1H), 7.77 (dd, *J* = 7.5, 2.0 Hz, 2H), 7.37–7.30 (m, 2H), 6.88 (dt, *J* = 8.3, 0.9, 0.9 Hz, 1H), 3.94 (s, 3H). ^13^C NMR (75 MHz, CD_3_OD) δ (ppm) = 189.9, (Cq), 154.8 (CH), 152.7 (Cq), 150.6 (Cq), 149.7 (Cq), 146.7 (Cq), 131.1 (CH), 129.0 (Cq), 125.7 (Cq), 124.6 (CH), 118.6 (Cq), 117.4 (CH), 116.0 (CH), 112.6 (CH), 110.6 (Cq), 108.8 (Cq), 57.0 (CH_3_). HRMS ESI (3000V): calculated for C_17_H_13_NO_5_Br (M+H^+^) 389.9972, found 389.9964.

#### (E)-3-(3-fluoro-4-hydroxy-5-methoxyphenyl)-2-(3,4-dihydroxybenzoyl)acrylonitrile (**10j**)

According to general procedure A, the reaction between malonitrile **8** (20.0 mg, 0.075 mmol) and 5-fluorovanilin (12.8 mg, 0.075 mmol) afforded the crude product that was used without purification in the deprotection according to general procedure B. This afforded **10j** (14 mg, 56%) as an orange solid. ^1^H NMR (300 MHz, DMSO) δ (ppm) = 10.66 (bs, 1H), 10.08 (bs, 1H), 9.54 (bs, 1H), 7.97 (s, 1H), 7.70–7.60 (m, 2H), 7.30–7.22 (m, 2H), 6.92–6.83 (m, 1H), 3.86 (s, 3H). ^13^C NMR (75 MHz, DMSO) δ (ppm) = 187.5 (Cq), 153.2 (CH), 152.3 (Cq), 151.2 (Cq), 149.4 (Cq), 149.3 (Cq), 149.2 (Cq), 145.3 (Cq), 126.8 (Cq), 122.9 (CH), 117.6 (Cq), 116.3 (CH), 115.2 (CH), 111.7 (d, *J*C-F = 20.2 Hz, CF), 110.7 (CH), 99.4 (CH), 56.2 (CH_3_). ^19^F NMR (282 MHz, DMSO) δ (ppm) = –76.9. HRMS ESI (3000V): calculated for C_17_H_13_NO_5_F (M+H^+^) 330.0772, found 330.0764.

#### (E)-3-(3,4-dihydroxy-5-iodophenyl)-2-(3,4-dihydroxybenzoyl)acrylonitrile (**10k**)

According to general procedure A, the reaction between malonitrile **8** (30.0 mg, 0.113 mmol) and 3,4-bis(methoxymethoxy)-5-iodobenzaldehyde (39.8 mg, 0.113 mmol) afforded the crude product that was used without purification in the deprotection according to general procedure B. This afforded 10k (11 mg, 23%) as a yellow solid. ^1^H NMR (300 MHz, CD_3_OD) δ (ppm) = 7.82 (d, J = 2.0 Hz, 1H), 7.78 (s, 1H), 7.69 (d, J = 2.1 Hz, 1H), 7.36–7.27 (m, 2H), 6.92–6.84 (m, 1H). ^13^C NMR (75 MHz, CD_3_OD) δ (ppm) = 190.3 (Cq), 154.8 (CH), 153.3 (Cq), 152.6 (Cq), 146.7 (Cq), 146.1 (Cq), 136.6 (CH), 129.1 (Cq), 126.7 (Cq), 124.5 (CH), 118.5 (Cq), 117.4 (CH), 116.4 (CH), 115.9 (CH), 107.8 (Cq), 84.2 (Cq). HRMS ESI (3000V): calculated for C_16_H_11_NO_5_I (M+H^+^) 423.9676, found 423.9688.

#### (E)-3-(4-hydroxy-3-iodo-5-methoxyphenyl)-1-phenylprop-2-en-1-one (**12**)

According to general procedure C, benzophenone (90.6 μL, 0.78 mmol) was reacted with 3-iodo-5-methoxy-4-(methoxymethoxy)benzaldehyde (250 mg, 0.78 mmol) and KOH (1.96 g, 35 mmol). This afforded the crude product that was purified using column chromatography (EtOAc:heptane / 0:100→ 40:60; gradient) to afford MOM-protected **12** (246 mg, 76%) as a yellow oil that solidified upon standing. 50 mg, 0.12mmol was directly used in the deprotection according to general procedure B. This afforded **12** (40 mg, 89%) as a yellow solid. ^1^H NMR (300 MHz, CD_3_OD) δ (ppm) = 8.05–7.95 (m, 2H), 7.70–7.45 (m, 5H), 7.37 (d, *J* = 16 Hz, 1H), 7.06 (d, *J* = 1.8 Hz, 1H), 6.46 (s, 1H), 3.95 (s, 3H). ^13^C NMR (75 MHz, CD_3_OD) δ (ppm) = 190.1 (Cq), 147.9 (Cq), 146.1 (Cq), 143.5 (CH), 138.1 (Cq), 132.7 (CH), 131.6 (CH), 129.0 (Cq), 128.6 (2xCH), 126.4 (2xcH), 120.8 (CH), 110.1 (CH), 81.6 Cq), 56.3 (CH_3_). HRMS ESI (3000V): calculated for C_16_H_14_IO_5_ (M+H^+^) 380.9982, found 380.9978

#### (E)-2-(hydroxy(phenyl)methyl)-3-(4-hydroxy-3-iodo-5-methoxyphenyl)acrylonitrile (**13**)

Compound **5a** (81 mg, 0.2 mmol) and CeCl_3_ (49 mg, 0.2 mmol) was suspended in MeOH (3 mL). NaBH_4_ (49 mg, 0.126 mmol) was added to the mixture at once at 0 °C. The mixture was allowed to stirr for 1.5 hours at 0 °C. Water (3 mL was carefully added to the mixture. The pH was adjusted to 4 using 1N HCl, This mixture was extracted with CH_2_Cl_2_ (3 x 10 mL), the organic layers were combined, dried over Na_2_SO_4_, filtered and concentrated *in vacuo*.Column chromatography (1:3 / heptane:EtOAc) afforded **13** as an off-white solid. ^1^H NMR (300 MHz, CHCl_3_) δ (ppm) = 7.61 (d, *J* = 1.9 Hz, 1H), 7.48–7.31 (m, 6H), 7.10 (s, 1H), 6.49 (s, 1H), 5.43 (s, 1H), 3.90 (s, 3H). ^13^C NMR (75 MHz, CDCl_3_) δ (ppm) = 147.9 (Cq), 145.9 (Cq), 141.3 (CH), 139.8 (Cq), 133.8 (CH), 128.9 (2xCH), 128.8 (CH), 127.0 (Cq), 126.4 (2xCH), 117.5 (Cq), 112.3 (Cq), 109.3 (CH), 80.8 (Cq), 75.5 (CH), 56.4 (CH_3_). HRMS ESI (3000V): calculated for C_17_H_14_NO_3_I (M+H^+^) 408.0091, found 408.0090.

#### 3-hydroxy-2-(4-hydroxy-3-iodo-5-methoxybenzyl)-3-phenylpropanenitrile (**14**)

Compound **5a** (81mg, 0.2mmol) was dissolved in MeOH (3 mL). NaBH_4_ (4.7 mg, 0.125 mmol) was added at once at 0 °C. The mixture was warmed to r.t. and stirred for 1 hour. Water (3 mL) was carefully added to the mixture. The pH was adjusted to 4 using 1N HCl and the mixture was subsequently extracted with CH_2_Cl_2_ (2 x 10 mL). Preparative LCMS afforded a 2:1 mixture of diastereomers **14** (18 mg, 22%) as a white solid. ^1^H NMR (300 MHz, CDCl_3_, 2:1 mixture of diastereomers) δ (ppm) = 7.34–7.19 (m, 10H_Ar_ D_major_/D_minor_) 7.12 (s, 1H, D_major_), 7.02 (d, *J* = 1.8 Hz, 1H, D_minor_). 6.97 d, *J* = 1.8 Hz, D_major_), 6.63 (d, *J* = 1.8 Hz, 1H, D_minor_), 6.61 (d, *J* = 1.8 Hz, D_major_), 5.94 (bs, 2H, D_major_/D_minor_), 4.68 (d, *J* = 5.1 Hz, 1H, D_minor_), 4.66 (d, *J* = 5.1 Hz, 1H, D_major_), 3.74 (s, 3H, D_major_), 3.73 (s, 3H, D_minor_), 2.99 (ddd, *J* = 9.6, 6.8, 4.5 Hz, 1H, D_minor_), 2.88 (ddd, *J* = 9.2, 6.0, 5.1 Hz, 1H, D_major_), 2.8–2.57 (m, 4H, D_major_/D_minor_). ^13^C NMR (75 MHz, CDCl_3_, 2:1 mixture of diastereomers) δ (ppm) = 146.0 (Cq, D_major_), 146.0 (Cq, D_minor_), 145.0 (Cq, D_major_), 145.0 Cq, D_minor_), 140.0 (Cq, D_major_), 139.7 (Cq, D_minor_), 130.7 (CH, D_minor_), 130.6 (CH, D_major_), 130.3 (Cq, D_major_), 130.2 (Cq, D_minor_), 129.1 (CH, D_minor_), 129.0 (CH, D_major_), 129.0 (2xCH, D_major_), 128.9 (2xCH, D_minor_), 126.4 (2xCH, D_minor_), 126.1 (2xCH, D_major_), 119.3 (Cq, D_major_), 119.2 (Cq, D_minor_), 111.9 (CH, D_minor_), 111.7 (CH, D_major_), 81.3 (Cq, Dmajor), 81.2 (Cq, D_minor_), 73.2 (CH, D_minor_), 73.1 (CH, D_major_), 56.3 (CH_3_, D_major_), 56.3 CH_3_, D_minor_), 43.2 (CH, D_major_), 42.6 (CH, D_minor_), 34.5 (CH_2_, D_major_), 33.2 (CH_2_, D_minor_). HRMS ESI (3000V): calculated for C_17_H_17_NO_5_I (M+H^+^) 410.0148, found 410.0153.

### Enzyme Activity Assays

#### Assay reagents

Pup_(33–63)_-Glu(AMC) used in this study was prepared according to the literature procedure.^27^ All other reagents were obtained from commercial suppliers.

#### Dop, FabD∼Pup, Ino1∼Pup, and PafA purification

*M. smegmatis* Dop, *Mtb* PafA-His_6_ and pupylated *Mtb* FabD and Mtb Ino1 were purified as described elsewhere.^21^

#### Data analysis

Data analysis and curve fitting was performed using GraphPad Prism V.7 (GraphPad Software, La Jolla, CA).

### Screening setup

The compounds were acoustically dispensed into black 384-well non-binding polystyrene microplates (Corning 3820) using an Echo 550 (Labcyte inc., Sunnyvale, CA). For the primary screening 15 nL of 10 mM DMSO solutions were dispensed per well (1% final DMSO concentration). Subsequently, buffer (5 μL; TRIS (50 mM, pH = 8.0), NaCl (50 mM), MgCl (20 mM), cysteine (1 mM), ATP(1 mM) and CHAPS (1 mg/mL)); Dop in buffer (5 μL, 30 nM) followed by 5 μL substrate in buffer (5 μL, 750 nM) after 30 min incubation were added via a MultiflowFX dispenser (Biotek, Winooski, VT). Single point fluorescence of the reactions was measured after 60 minutes using a CLARIOstar plate reader (BMG Labtech, Ortenberg, Germany).

### IC_50_ determinations

#### Assay buffer

All reactions were performed in freshly prepared assay buffer at room temperature. The buffer contained TRIS (50 mM, pH = 8.0), NaCl (50 mM), MgCl (20 mM), cysteine (1 mM), ATP (1 mM, unless otherwise indicated) and CHAPS (1 mg/mL).

For all the compounds stock solutions in DMSO (10 mM) were prepared and 10-point three-fold serial dilutions in buffer were prepared with a top concentration of 300 μM (100 μM final higest conc.). 10 μL of each concentration was transferred to a black 384-well non-binding polystyrene microplates (Corning 3820). 10 μL of a solution of Dop in buffer (30 nM; 10 nM final conc.) was added after which the reactions were incubated 30 minutes in the dark. The depupylation reaction was initiated by addition of 10 μL of substrate in buffer (750 nM; 250 nM final conc.). The reactions were monitored by measuring the increase of fluorescence emission at 450 nM (λex = 360 nm) that correlates with hydrolysis of AMC from the substrate using a CLARIOstar plate reader (BMG Labtech). The fluorescence signal was measured every minute for 1 hour. The data was fitted and IC_50_ values were derived using GraphPad Prism V.7 (GraphPad Software, La Jolla, CA). All data presented as mean ± s.d. (n = 3).

### Mode of inhibition study

10-point three-fold serial dilutions of **1** in buffer were prepared with a top concentration of 100 μM. Separate series were incubated with Dop (10 nM) for 5 and 60 minutes. Directly after the addition of the substrate, the course of the reactios was monitored similarly to the IC_50_ measurements..

#### Specificity assay

10-point three-fold serial dilutions of **1** in buffer were prepared with a top concentration of 300 μM (100 μM final conc.). Separate series were incubated with regular Dop concentration (30 nM; 10 nM final conc.) and with 10-fold concentration Dop (300 nM; 100 nM final conc.) for 3 minutes. The assay was performed using the procedure for determination of the IC_50_ values.

#### Jump-dilution assay

Dop was pre-incubated for 30 minutes with 10x the inhibitor IC_50_ value followed by 100-fold dilution in buffer. The fluorescence increase was followed for 20 minutes as described above.

### ATP/ADP-competition assay

#### Determination of the Km for ATP

10-point doubling dilutions of ATP in buffer were prepared with a top concentration of 48 mM (16 mM final conc). These solutions were 30 min incubated with Dop (30 nM, 10 nM final conc.). Pup_(33–63)_-Glu(AMC) (750 nM; 250 nM final conc.) was added and Dop activity was followed for 20 minuted as describes above. The initial rates were fitted in the Michaelis Menten equation and the K_m_ and V_max_ values were derived using GraphPad Prism V.7 (GraphPad Software, La Jolla, CA).

### In vitro assays on native substrates

**Buffer A**: 50 mM Tris, pH 8; 10% glycerol; 20 mM MgCl_2;_ 150 mM NaCl; 5 mM ATP; 1 mM DTT. **Buffer B**: 50 mM NaH_2_PO_4_, pH 8; 10% glycerol; 20 mM MgCl_2;_ 150 mM NaCl; 5 mM ADP; 1 mM DTT.

#### Dop depupylation assays

10 nM Dop was incubated with the indicated amount of compound in Buffer A for 30 min at room temperature. The substrate Myc-PupE∼FabD-His_6_ was then added to a final concentration of 0.25 µM in a final volume of 50 µl. After 30 minutes, 20 µl of sample was withdrawn and added to 10 µl of 3X SDS sample buffer. Samples were boiled for 10 min before analysis by 10% SDS-PAGE gels (NuPAGE™ 10% Bis-Tris Protein Gels, 1.0 mm, Thermo-Fisher Scientific, Catalog # NP0301BOX). For immunoblot analysis, samples were transferred to a nitrocellulose membrane, blocked with 3% bovine serum albumin (BSA), incubated with primary polyclonal antibodies to *Mtb* FabD-His_6_ in 3% BSA/PBS, and then incubated with green fluorfescent IRDye 800CW goat anti-rabbit IgG (H+L) (926-32211, Li-COR) in 3% BSA/PBS. Detection of the fluorescent signal was performed with a Li-Cor Biosciences Odyssey CLx imaging system.

#### PafA depupylation assays

50 nM PafA was incubated with the indicated amount of compound in Buffer B for 30 min at room temperature. The substrate Myc-Pup∼FabD-His_6_ or Pup∼Ino1∼His_6_ was then added to a final concentration of 0.25 µM in a final volume of 50 µl. After 2 hours, 30 µl of sample was withdrawn and added to 10 µl of 4X SDS sample buffer. Samples were boiled for 10 min before analysis by 10% SDS-PAGE gels and immunoblotting with primary polyclonal antibodies to FabD or Ino1 in 3% BSA/TBST, and then incubated with horseradish peroxidase (HRP) conjugated anti-rabbit secondary antibodies (Thermo-Fisher Scientific) in 3% BSA/TBST. Detection of HRP was performed using SuperSignal West Pico.

#### PafA pupylation assays

50 nM PafA was incubated with the indicated amount of compound in Buffer A and 0.25 µM FabD-his for 30 min at room temperature. PupE was then added to a final concentration of 4 µM in a final volume of 50 µl. After 6 hours, 30 µl of sample was withdrawn and added to 10 µl of 4X SDS sample buffer. Samples were boiled for 10 min before analysis by 10% SDS-PAGE gels and immunoblotting with primary polyclonal antibodies to FabD in 3% BSA/TBST as above.

## ASSOCIATED CONTENT

### Supporting Information

Supplementary figures S1–S3 (PDF)

## AUTHOR INFORMATION

### Author Contributions

The manuscript was written through contributions of all authors. All authors have given approval to the final version of the manuscript.

### Notes

The authors declare no competing financial interest.

## ABBREVIATIONS

Tb: tuberculosis;
*Mtb*: *mycobacterium tuberculosis*;
PPS: Pup proteasome system;
Dop: deamidase op Pup;
PafA: proteasome accessory factor A;
AMC: 7-amino-4-methylcoumarin;
FabD: Malonyl CoA-acyl carrier protein transacylase;
Ino1: Inositol-3-phosphate synthase.

