## Supplementary Figures 1-3 for "Discovery and Optimization of Inhibitors for the Pup Proteasome System in *Mycobacterium tuberculosis*"

### Contents

**Supplementary figure 1** (Page S2)
**Supplementary figure 2** (Page S2)
**Supplementary figure 3** (Page S3)

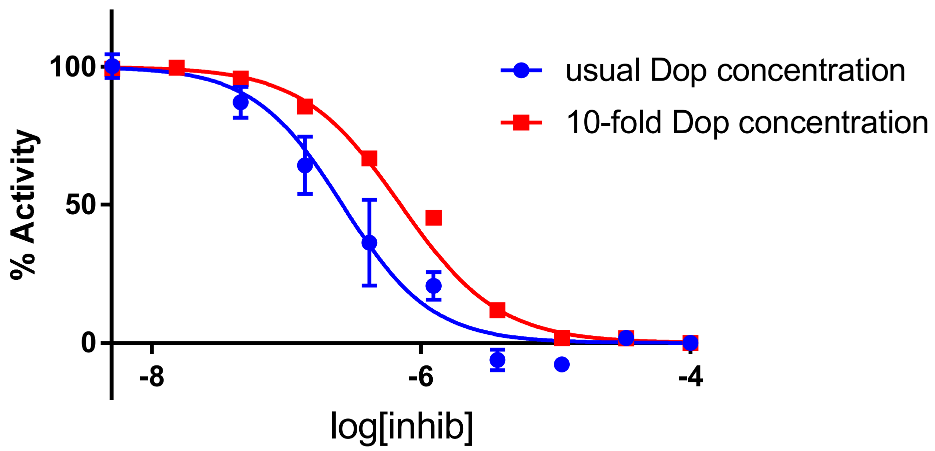


**Supplementary Figure 1**. $IC_{50}$ curves for the 'ratio test'. $IC_{50}$ value normal Dop concentration = 0.26±0.06 µM; $IC_{50}$ value 10-fold Dop concentration = 0.70±0.03 µM. The reported values are the means of one experiment ± SD run in triplicate. Because the deviation between the values for the 10-fold Dop concentration were very small, the error bars are not omitted.

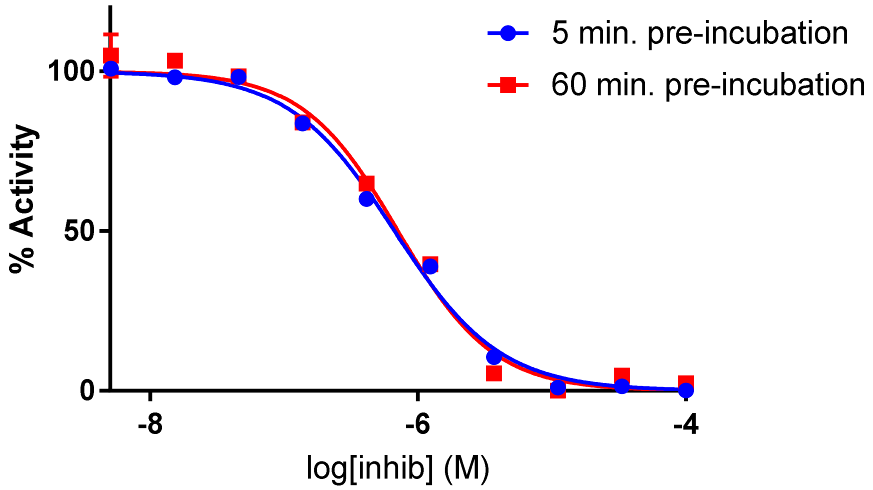


**Supplementary Figure 2**. $IC_{50}$ curves of compound **1** with 5 and 60 minutes inhibitor/enzyme pre-incubation. $IC_{50}$ value for 5 min preincubation = 0.47±0.05 µM; $IC_{50}$ value for 60 min preincubation = 0.71±0.08 µM. The reported values are the means of one experiment ± SD run in triplicate.

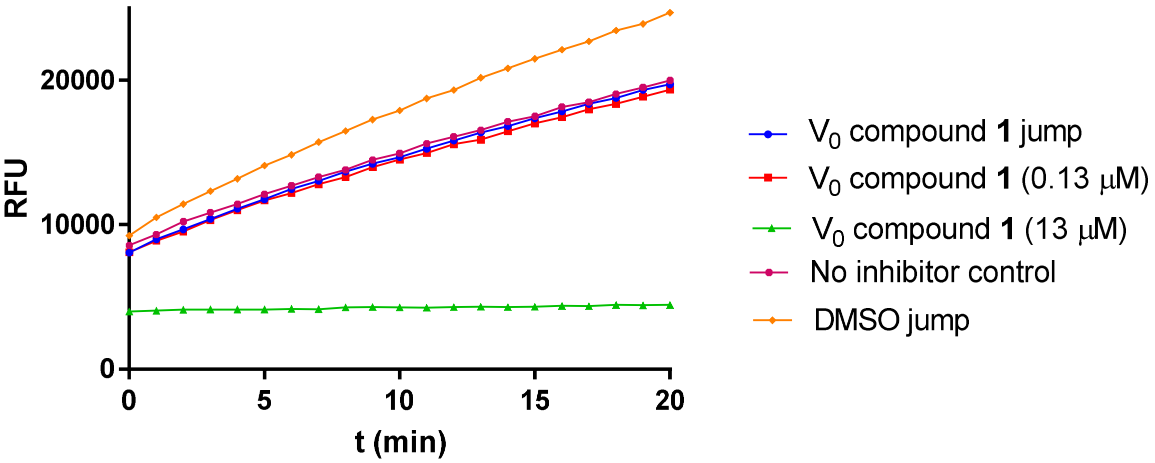


**Supplementary Figure 3**. Reaction curves for the jump dilution experiment with compound **1**. RFU = relative fluorescence units. [Dop] before jump dilution = 100 nM; [Dop] after jump dilution = 10 nM.
